# Low-Toxin *Clostridioides difficile* RT027 Strains Exhibit Robust Virulence

**DOI:** 10.1101/2022.03.18.484943

**Authors:** Farhan Anwar, Bryan Angelo P. Roxas, Kareem W. Shehab, Neil Ampel, VK Viswanathan, Gayatri Vedantam

**Author notes:** Correspondence to: Gayatri Vedantam, Ph.D., University of Arizona, School of Animal & Comparative Biomedical Sciences, 1117 E. Lowell St. Bldg. 90, Tucson, AZ 85721.

## Abstract

*Clostridioides difficile* is a leading cause of healthcare-associated infections worldwide. Currently, there is lack of consensus on a single optimal diagnostic method for *C. difficile* infection (CDI). Nucleic acid amplification tests (NAAT) that detect toxin genes are highly sensitive, but their specificity limitations could inflate CDI rates. Alternate multi-step diagnostic algorithms emphasize the detection of *C. difficile* toxins TcdA/TcdB, and are premised on the rationale that stool toxin-negative (Tox^-^) CDI patients have less severe disease, shorter diarrhea duration, and fewer disease complications. There have been no systematic assessments, however, of the virulence of *C. difficile* strains from Tox^-^/NAAT^+^ (discrepant) specimens. In our prospective analysis of 1243 *C. difficile-*positive patient stool specimens from three Southern Arizona hospitals, 31% were discrepant. Ribotype 027 (RT027) strains were recovered from 221 specimens; of these, 23% were discrepant. Post-culture, RT027 strains produced a range of toxin amounts including levels lower than that of the non-epidemic strain CD630. These low-toxin RT027 (LT-027) strains harbored both *tcdA* and *tcdB* genes, and their culture supernatants were cytotoxic to cultured fibroblasts. We confirmed robust colonization and virulence of a subset of LT-027 strains using multiple rodent models; lethality in animals infected with LT-027 strains was comparable to that potentiated by a high-toxin RT027 strain. Comparative genomics and proteomics analyses of several LT-027 strains identified unique genes and altered protein abundances relative to closely-related high-toxin strains. Collectively our data highlight the robust virulence and clinical relevance of low-toxin-producing, RT027 *C. difficile* isolates.

## INTRODUCTION

*Clostridioides difficile* (CD), an anaerobic, spore-forming, Gram-positive bacterium, is the leading cause of healthcare-associated infections in the United States and Europe [1]. In the US, over 225,000 cases of *C. difficile* infection (CDI) occur annually [2], with associated total medical costs ranging from $1.9-7 billion [3]. The past 20 years have seen the emergence of outbreak-associated strains, particularly those belonging to the ribotype 027 [RT027; also called (REA) type BI or North American Pulsed-Field (NAP) type NAP1]. These strains persist in clinical settings and have been associated with severe disease [4].

*C. difficile* strains may produce up to three toxins: TcdA (Toxin A), TcdB (Toxin B), and Binary Toxin (CDT) [5]. TcdA and TcdB, encoded within a pathogenicity locus (PaLoc), are glucosyltransferases that target host-cell GTPases including Rho, Rac and Cdc42 [6]. The resulting inhibition of these small GTPases contributes to cytoskeletal changes, epithelial barrier disruption and, on prolonged exposure, host cell death [5]. Non-toxigenic *C. difficile* strains lack the entire PaLoc region [7].

Rapid and accurate CDI diagnosis can guide treatment, limit inappropriate antibiotic use, and curtail transmission. There is, however, considerable variability in published CDI diagnosis guidelines [8]. Broadly, diagnostic protocols rely on three assays: nucleic acid amplification tests (NAATs) that detect toxigenic *C. difficile*, enzyme immunoassays (EIA) for glutamate dehydrogenase (GDH), an enzyme expressed in all strains (including non-toxigenic *C. difficile*), and EIA for TcdA and TcdB [9]. Some experts advocate for sole use of the highly sensitive NAAT approach, but others have argued that this test fails to distinguish between ‘true’ CDI, and non-CDI diarrhea with concomitant *C. difficile* colonization [8]. Alternate “two-step” protocols thus utilize NAAT or GDH EIA to determine the presence of *C. difficile*, coupled with the detection of *C. difficile* toxins by toxin EIA. CDI is deemed to be highly unlikely if free toxins are not detected in the stool (Tox^-^). However, since CDI cannot be ruled out when clinical test results are discrepant (Tox^-^ /NAAT^+^), the decision to treat for CDI in these instances is based on clinical judgement [1,10].

While the diagnostic implications of various testing strategies have been extensively debated, the factors contributing to discrepant laboratory results have hitherto not been systematically addressed [11,12]. A Tox^-^/NAAT^+^ finding may result from limitations of the specific diagnostic tests or clinical laboratory protocols used; this can sometimes be resolved by re-testing, which is not recommended [9]. Alternatively, the test results may reflect a discrepancy: patients may harbor toxigenic *C. difficile* but various factors, including decreased pathogen abundance or host impacts, may depress toxin abundance in the stool. Additionally, some *C. difficile* isolates may produce extremely low levels of toxin; indeed, it is well-recognized that toxigenic *C. difficile* strains, including those belonging to RT027, are highly diverse in the amounts of toxins they produce [13]. Strains that consistently produce very low amounts of toxins have not been specifically explored, and their virulence potential has not been established in animal models.

In our survey of three local hospitals, we found a substantial number of discrepant Tox^-^/NAAT^+^ stool specimens (31%). After verifying the presence of toxigenic *C. difficile* strains in these specimens, we identified RT027 strains that consistently produced very low amounts of TcdA/TcdB (LT-027 strains). We assessed the virulence and colonization of several LT-027 strains using the Golden Syrian hamster model of acute infection, and a mouse colonization model. Several LT-027 strains were characterized via whole genome sequencing (WGS) as well as comparative, quantitative proteomics. Our observations have implications not only for CDI diagnosis and clinical management, but also for *C. difficile* virulence.

## METHODS

### Specimen acquisition and recovery of C. difficile

Stool specimens from patients suspected to have CDI were obtained over a period of 7 years from three hospitals in Tucson, Arizona. All specimens were designated “to-be-discarded” and were de-identified prior to acquisition. This study was deemed not to constitute human subjects research and was exempt from full Institutional Review Board approval.

An aliquot of the stool specimen was plated to isolation on taurocholate-cefoxitin-cycloserine-fructose agar (TCCFA) which is selective for *C. difficile*. PCR amplification of *tcdB* was used to confirm presence of toxigenic *C. difficile*. Supplemental Table S1 lists primers and strains used for this study. For the studies described below, and unless otherwise indicated, isolates were grown in Brain-Heart Infusion broth (Bacto BHI; BD, Franklin Lakes, New Jersey), and cultures maintained in 25% glycerol at -80°C for long-term storage.

### Spore Preparation

*C. difficile* overnight cultures were sub-cultured in 50 mL BHI broth for 14 days, harvested by centrifugation, and washed three times with phosphate-buffered saline (PBS). The pellets were re-suspended in sterile water, and remaining vegetative bacteria were killed by heat-shock at 65°C for 15 minutes. Spores were purified via at least three washes in distilled water, or via ficoll gradient centrifugation as previously described, and stored in water at 4°C [14].

### Ribotyping and Colony PCR

Overnight BHI broth cultures of *C. difficile* strains were centrifuged and resuspended in 1 mL of Tris-EDTA (pH 7) and processed for genomic DNA extraction and ribotyping PCR as per published protocols (Supplemental Table S1) [15,16]. PCR products were separated via capillary electrophoresis, and the ribotype established via comparison to the online database Webribo [17].

The presence of toxin genes was verified via PCR amplification of sequences specific to *tcdA, tcdB* and binary toxin genes (Supplementary Table S1), respectively, as previously described [16,18]. Isolated *C. difficile* colonies on TCCFA were transferred to 25 µL of PCR-Lyse™ (Epicentre, Madison, WI, USA) solution, incubated at 99°C for 5 minutes, and immediately placed on ice. The sample was centrifuged, and the supernatant was used for PCR analysis.

### Toxin Analysis

Clinical and reference *C. difficile* strains were grown in 5 ml BHI broth for 72 hours. The cultures were then centrifuged, and cell-free supernatants were clarified using 0.22 µm filters (Argos Technologies, Elgin, IL, USA). Toxin levels in 50 μL of the supernatants was assessed using the Techlab® *C. difficile* Tox A/B II™ EIA kit (Techlab, Blacksburg, VA, USA) as per manufacturer’s instructions, and optical density measured at 450 nm. Total protein concentration was assessed using the Pierce™ BCA Protein Assay Kit (ThermoFisher Scientific, Waltham, MA, USA), and toxin levels reported as OD450nm/mg total protein. All data were collected in biological triplicate. Immunoblotting was also performed on select isolates. Following overnight growth and sub-culture in 50 mL BHI broth for 72 hours, the cultures were centrifuged, and 30 mL of the supernatant was concentrated using a 100 kDa filter (Amicon Ultra, Millipore). 50 μg of protein from the concentrated supernatants were used for immunoblot analysis.

### Cytotoxicity Assays

Cultured Vero fibroblasts were propagated according to previously published protocols^22^ in 96-well plates with Eagle’s Minimum Essential Media (EMEM; Corning Life Sciences, Corning, NY, USA) containing 10% fetal bovine serum (FBS; Corning Life Sciences, Corning, NY, USA). Clarified *C. difficile* supernatants (described above) sequentially diluted in EMEM + FBS were added to each well and incubated for 24 hours at 37°C with 5% CO2. Cells were visualized using a bright field microscope (Olympus BHTU; Olympus, Center Valley, PA, USA) after specific time intervals, and the degree of cell rounding was scored on a scale of 1 (little-to-no cell rounding) to 4 (complete cell rounding). TcdA/TcdB-dependent rounding was verified via neutralization with anti-TcdA and anti-TcdB monoclonal antibodies (Abcam, Cambridge, MA, USA). A minimum of 10 fields per treatment, with three independent replicates, were assessed.

### Antibiotic Susceptibility Assay

Susceptibility of select *C. difficile* isolates to cefotaxime, rifampicin, levofloxacin, metronidazole and vancomycin was determined using Clinical and Laboratory Standards Institute (CLSI) protocols (M11-A8) [19]. Pure cultures were grown overnight, sub-cultured to 0.5 McFarland standard, and plated on Brucella agar with hemin, vitamin K1, and 5% horse blood (BD, Franklin Lakes, New Jersey). Antibiotic susceptibility was determined using the bioMérieux Etest® platform (bioMérieux, Durham, NC, USA).

### Golden Syrian Hamster Infections

Golden Syrian hamsters were used to assess virulence of select *C. difficile* strains [20,21]. Seven-to-eight-week-old male hamsters (90-100 grams; Charles River, Wilmington, MA, USA) were orally administered clindamycin (30 mg/kg) (Pfizer, New York, NY, USA) 72 hours prior to infection. For pilot studies, animals were orally infected with 100 spores of *C. difficile* LT-027 strain (GV106, GV135, or GV148) or a high-toxin RT027 strain (BI-1) [22]. For fully-powered studies, animals were infected with 100 spores of the LT-027 strain GV148 (N = 8), BI-1 (N = 6), or a non-toxigenic strain (T7; N = 2). In both studies, animals were monitored for overt disease every 6-12 hours (diarrhea, inappetence, ruffled fur, lethargy). Surviving animals were humanely euthanized at the end of the study using a commercial euthanasia solution (270 mg/kg sodium pentobarbital: Merck, Kenilworth, NJ, USA). *C. difficile* burden in stool and cecal contents was determined by plating on TCCFA. Cecal toxin burden was determined as detailed above.

#### Mouse Infections

Intestinal colonization of select *C. difficile* strains was assessed using a non-lethal mouse model of infection [23]. 10-week-old male C57BL/6 mice (Jackson Laboratories, Bar Harbor, ME, USA) were pre-treated with cefoperazone (0.5 mg/mL) in the drinking water for 10 days, followed by intraperitoneal clindamycin (10 mg/kg) (Pfizer, New York, NY, USA) on day -1. The animals were administered 10^6^ spores of BI-1, GV106, GV135, or GV148, via oral gavage on day 0. Animals were weighed and monitored for signs of stress daily. Surviving animals were humanely euthanized at the end of the study using a commercial euthanasia solution described above. Stool pellets or cecal content were collected daily, resuspended, homogenized in PBS, serially diluted, and plated onto TCCFA for isolation and bacterial burden (CFU/g of stool or cecal tissue).

### Serum Collection and Immunoblots of Surface Proteins

Mice infected with high toxin strains (BI-1, GV599) or LT-027 strains (GV148, GV1002) were humanely euthanized on day 14 post-infection, and blood was collected via cardiac puncture. Serum was isolated by centrifuging the blood for 15 minutes at 400 rpm. HALT protease (ThermoFisher Scientific, Waltham, MA, USA) was added to the serum and stored at -80°C.

*C. difficile* surface proteins were extracted for immunoblotting as previously reported [14]. Briefly, each strain was grown overnight in 50 mL of BHI media. The cultures were centrifuged, and the pellets were resuspended in 500µL of 0.2M glycine (pH2.2). 30µg of surface extract was separated on a 4-20% gradient acrylamide gel and transferred to a PVDF membrane. These were probed with collected serum at a 1:500 dilution overnight, and an HRP-conjugated goat anti-mouse secondary antibody (1:10,000 dilution; Abcam, Cambridge, MA, USA) for 1 hour; the blots were developed using SuperSignal West Femto chemiluminescent substrate (ThermoFisher Scientific, Waltham, MA, USA) and imaged with a Chemi-doc Imager (Bio-Rad, Hercules, CA, USA).

### Comparative Genomics and Proteomics

Whole genome sequencing of 15 LT-027 and 3 RT027 isolates was performed using the Illumina MiSeq platform at the University of Arizona Genomics Core. The sequences were assembled using the CLC Genomics Workbench (Qiagen, Hilden, Germany) and annotated using RAST (NMDPR, Chicago, Illinois) (Supplemental Table S2). Alignment-free composition vector tree analysis of these genomes, and 10 publicly-available RT027 genomes, was performed using CVTree 3.0 [24] and visualized using the Interactive Tree of Life v4 (ITOL; https://itol.embl.de/) [25]. Eleven LT-027 strains were compared to previously published and publicly available RT027 genomes (N = 10). Similarly, 5 discrepant LT-106 strains and 4 high-toxin RT106 strains were sequenced, and composite vector trees were created.

For comparative proteomic analysis, strains were grown in BHI media overnight, and then sub-cultured 1:50 in BHI. Samples were collected at mid-logarithmic phase and processed for iTRAQ proteomic analysis as previously reported [26,27]. The *C. difficile* strain BI-1 protein database was used as the reference. Differentially-abundant proteins were identified via multiple stringent statistical tests as described below.

### Statistical analyses

XLSTAT was used for statistical analysis and generating PCA plots. For bacterial burden, Student’s t-tests and ANOVA was used to determine differences between strains. For studies with large number of observations for various time points (such as cytotoxicity analyses), goodness-of-fit statistical analyses were performed using ANOVA followed by stringent Tukey tests. *p* values of <0.05 were considered significant.

For proteomics, additional tests established significance of protein abundance changes. Specifically, three parameters had to be satisfied prior to a protein being categorized as differentially abundant [26]: (1) a (stringent) false discovery rate (FDR) of 1%, (2) a fold change cutoff determined by using a technical replicate in each experiment; and (3) hypergeometric testing, similar to the Fisher’s exact one-tailed test, where, in addition to the parameters described above, a significance of p<0.05 has to be achieved for differentially abundant proteins to be classified as such.

For all animal studies, χ2 analyses determined the percentage of animals colonized with *C. difficile*, the time interval between *C. difficile* challenge and colonization, and time between *C. difficile* challenge and death. Additionally, Kaplan–Meier survival curves were generated, followed by Log-Rank tests for post hoc analyses. Significance was determined by ANOVA, where goodness of fit and standardized coefficients of variance were derived.

#### Ethics Statement

All *in vivo* studies were carried out in strict accordance with the recommendations in the Guide for the Care and Use of Laboratory Animals of the National Institutes of Health. All animal studies were approved by the Institutional Animal Care and Use Committee of the University of Arizona (Protocol Number GV 14-526; PHS Assurance Number A3248-01; USDA Registration Number 86-R-0003). For human stool specimens, all samples were designated “to-be-discarded” and were de-identified before acquisition; this study was performed under an Institutional Review Board (IRB) Non-Human Subjects Research Data Use Committee Approval NRDUC # 1707612129.

## RESULTS

### *C. difficile* isolates from discrepant specimens express low levels of glucosylating toxins

The last systematic surveillance of *C. difficile* infections in Arizona was conducted in 1994 [28], prior to the emergence of current epidemic *C. difficile* strains. We conducted two independent CDI surveillance studies in Tucson-area hospitals. In a “snapshot” surveillance performed over 6 months in 2012, we obtained 93 independent *C. difficile* isolates upon selective culture of diarrheic stool specimens. Ribotyping of the 93 isolates revealed that the most frequent ribotype was RT027 (19.4%) (Figure 1A). In a subsequent prospective surveillance from 2015 to 2019, we isolated and ribotyped 1,150 *C. difficile* isolates. Of these, RT027 strains were the most numerous, accounting for 17.7% of all *C. difficile* isolated (Figure 1B). Thus, RT027 has been, and remains, the dominant *C. difficile* ribotype recovered in our hospital surveillance efforts. RT106, which represented only 3.2% of the isolates in 2012, is the second most common ribotype isolated (7%) which is consistent with broader trends across the United States [29,30].

**FIGURE 1:**
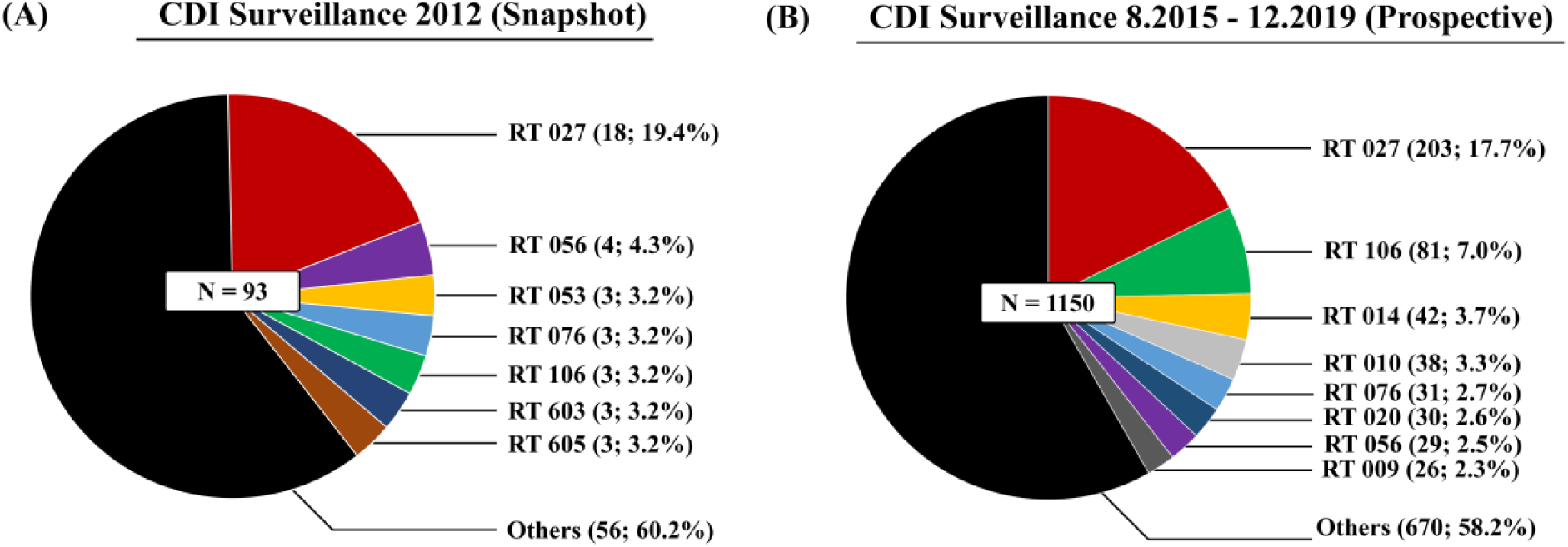
*Clostridioides difficile* Ribotype RT027 is Consistently Prevalent in Tucson-area Hospitals. The two charts depict ribotype distribution of *C. difficile* strains isolated from patient stool specimens collected from a ‘snapshot’ surveillance in 2012 (A; N = 93) and a prospective surveillance spanning between 2015-2019 (B; N = 1150). Ribotype frequency and percent of total sample size are shown in parenthesis (n, %). RT027 is the predominant ribotype for both periods. ‘Others’ refers to ribotypes that each constitute 2% or less of the total.

For the 2012 and 2015 – 2019 sampling periods, we were able to establish (by PCR) or infer (by ribotype association) that 80/93 and 781/1,150 specimens, respectively, harbored the toxin genes. Restricting analysis to the specimens for which clinical EIA results were available in the two periods, 54.5% (6/11) and 38.9% (384/988), respectively, were deemed to be ‘discrepant’ (Tox^-^/NAAT^+^). For all specimens harboring RT027 strains, 34% were discrepant.

To better appreciate the potential reasons for failing the toxin EIAs, we focused our studies on the first 93 samples obtained. The isolates were cultured in BHI for 72 hours and TcdA/TcdB abundance in 50 μL of cell-free supernatant was assessed via EIA (Figure 2A). The low-toxin producing strain CD630 and high-toxin-producing strain VPI 10463 were used as comparators. The 93 *C. difficile* strains produced highly divergent amounts of TcdA/TcdB. Toxin levels for 30/93 isolates (32.3%) were below the limit of detection (OD450 < 0.12). Including these, toxin levels of 74/93 isolates (79.5%) were at, or lower than, CD630. Notably, and consistent with other reports [13], there was substantial variability in toxin amounts produced by RT027 strains. Of the 18 recovered RT027 isolates, TcdA/TcdB levels of 13 (72%) were at, or below, that of CD630 (Figure 2A). Even after normalization for protein amounts, the 13 RT027 strains produced nearly ten-fold lower toxin levels relative to the reference, outbreak-associated comparator BI-1 (Figure 2B). Immunoblot analysis of select RT027 isolates confirmed decreased TcdA/TcdB expression compared to BI-1 (Figure 2C). We designated these RT027 strains that consistently produced >10-fold less toxin than BI-1 as “low-toxin 027” or “LT-027” strains. Importantly, LT-027 strains had similar growth curves as BI-1 (data not shown). These results suggest that, irrespective of ribotype, toxigenic *C. difficile* strains are highly divergent in the amounts of toxin they secrete. This could be one contributing factor to the Tox^-^/NAAT^+^ discrepancy in routine clinical diagnostic tests.

**FIGURE 2:**
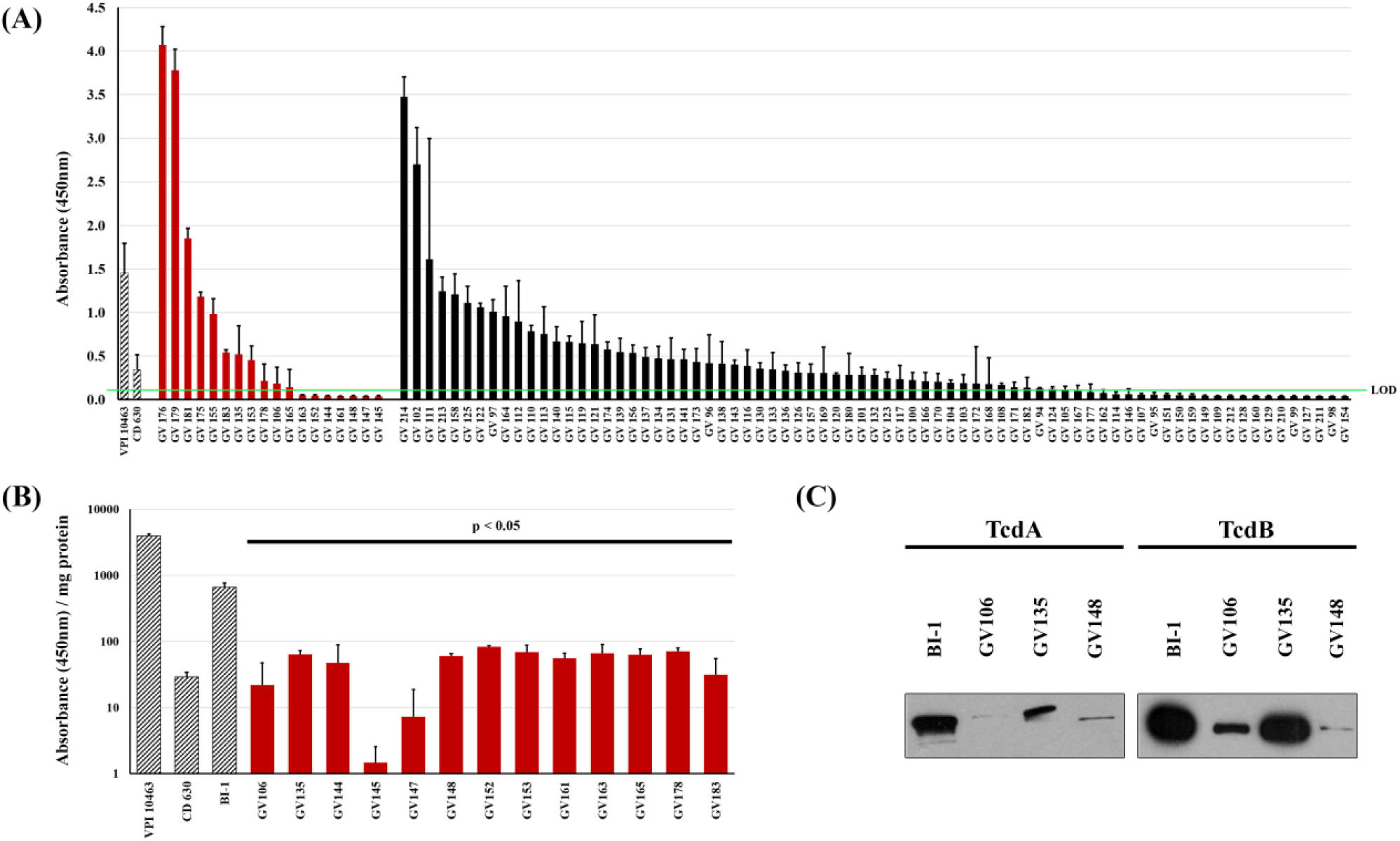
Clinical *C. difficile* strains display high variability in toxin production. TcdA/B ELISA assays of clarified stationary-phase supernatants from 93 *C. difficile* strains (A), normalized to volume, showed high variability in toxin abundance, even for RT027 strains (red bars; black bars include all other ribotypes). The high-toxin producer VPI 10463 (at 1:100 dilution) and low-toxin producer CD630 are included for comparison (hashed bars). Thirty strains registered readings below the limit of detection (LOD, green line = OD450 < 0.12). After normalization for protein amounts, supernatants of the thirteen RT027 strains shown in (A) had nearly ten-fold lower TcdA/B amounts relative to the comparator RT027 strain BI-1 (p<0.05; two sample, two-tailed t-test). Immunoblot analyses of concentrated supernatants (30 μg/lane) confirmed lower abundance of TcdA and TcdB in supernatants of three representative LT-027 strains relative to BI-1 (C).

### Systematic assessment of toxin production by Low-Toxin 027 *C. difficile isolates*

If the supposition that a threshold level of toxin activity is required to cause disease is correct, this could be achieved by secreting lower amounts of more potent toxins. We, therefore, used cytotoxicity assays to compare toxin activity. BI-1 induced Vero cell rounding at lower protein concentrations than the LT-027 strains GV106, GV135, GV148 (Figure 3A). When normalized to toxin amounts, however, LT-027 strains and BI-1 induced Vero cell rounding (Figure 3B) at similar rates. Collectively, these data suggest that while many clinical RT027 strains produce substantially less toxin than BI-1, those toxins are similarly cytotoxic to host cells.

**FIGURE 3:**
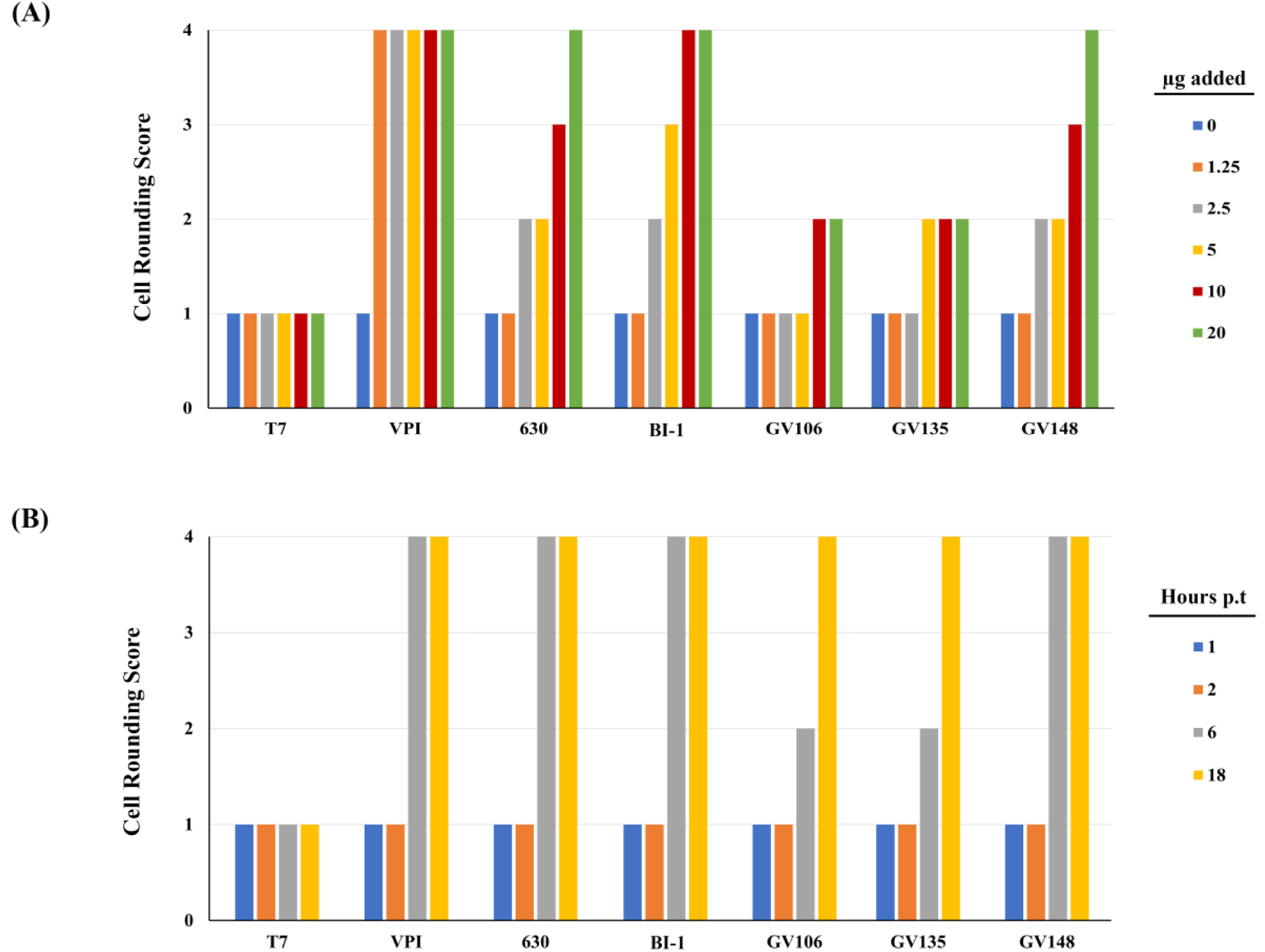
Cytotoxic activity of LT-027 strain supernatants is comparable to that of BI-1. Vero fibroblasts treated for 3 hours with filtered supernatants from BI-1 exhibited cell rounding at lower concentrations than supernatants of LT-027 strains GV106, GV135 and GV148 (A). After normalizing toxin amounts, LT-027 strains displayed comparable, or slightly slower, cytotoxicity kinetics as VPI 10463, BI-1, and CD630 (B). Averages were calculated from 10 fields per condition; triplicates of these sets resulted in the same average score and, therefore, no standard error is reported.

We next considered the possibility that distinct antibiotic resistance profiles for LT-027 strains, compared to historic RT027 strains, might confer a selective survival advantage *in vivo*, and account for higher bacterial burdens in vivo (and, thereby, increasing toxin amounts). The antibiotic susceptibility profile, as tested with the Etest® platform (Supplemental Table S3), revealed uniform susceptibility of all RT027 strains to vancomycin. While LT-027 strains exhibited variable susceptibility to other antibiotics, they did not collectively display a resistance pattern to distinguish them from BI-1.

### LT-027 *C. difficile* isolates cause lethal disease in hamsters

The correlation between stool toxin levels and symptomatic CDI continues to be actively debated [11,12,31]. This prompted us to assess if strains isolated from discrepant clinical samples, and that consistently produce low levels of toxin, can cause disease in the Golden Syrian hamster model of acute CDI. In a pilot study (N = 2), hamsters were infected with 100 spores of LT-027 isolates (GV106, GV135, or GV148), BI-1, or the non-toxigenic strain, T7. The LT-027 strains GV135 and GV148, like the reference BI-1 strain, caused 100% lethality within 24 hours of infection; LT-027 strain GV106 also induced 100% lethality, but with slightly delayed kinetics (60 hours post-infection) (Figure 4A). All the toxigenic strains induced hallmark CDI clinical signs including diarrhea, rapid weight loss, lethargy, inappetence and gross intestinal pathology. Cecal toxin abundance of TcdA/TcdB was, however, lower in LT-027 infected animals compared to BI-1-infected animals (Figure 4B). Thus, despite producing >10-fold lower level of toxins, LT-027 strains exhibited robust virulence in the hamster model of acute CDI, with disease progression indistinguishable from that of high-toxin (BI-1)-infected animals.

**FIGURE 4:**
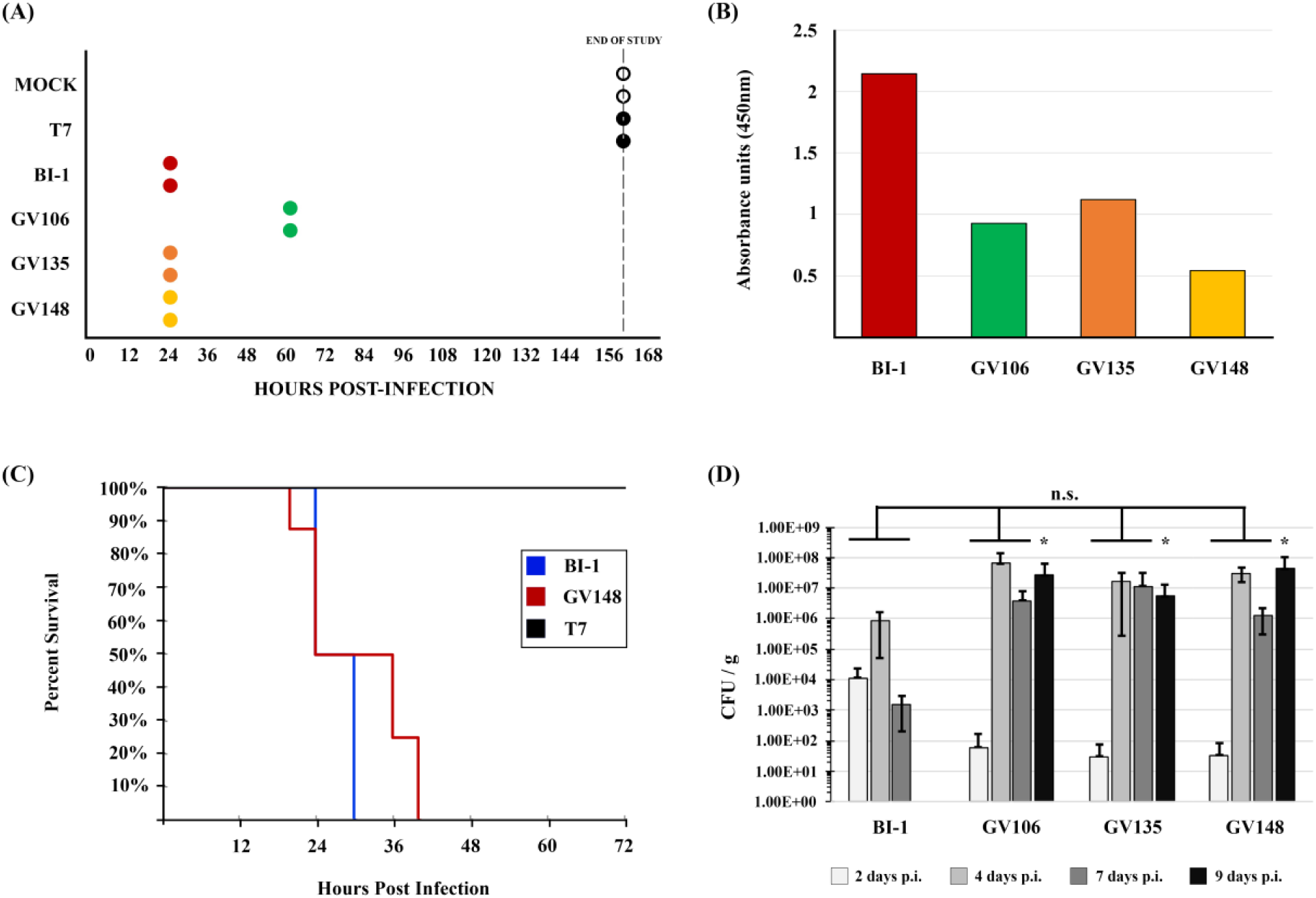
LT-027 Strains are Virulent and Exhibit a Colonization Advantage *in vivo*. In a pilot experiment, Golden Syrian hamsters infected with LT-027 strains succumbed to disease with a timeframe comparable to that of BI-1-infected animals (A; all groups N = 2). Animals infected with the non-toxigenic T7 strain did not display any symptoms and survived the infection. (B) Average cecal toxin levels (pilot, N = 2 / group) were lower in the LT-027-infected animals relative to BI-1, normalized to volume. (C) Kaplan-Meier survival curves of hamsters infected with the LT-027 strain GV148 (red, N = 8), BI-1 (blue, N = 6) and T7 (black, N = 2). There was no statistical difference for time-to-death between GV148 and BI-1 (log-rank, p-value > 0.05). Stool *C. difficile* burdens in C57BL/6 mice infected with select LT-027 strains increased more rapidly, and persisted for longer durations, relative to BI-1 (N = 5 for all conditions). There were no significant differences between each strain (independent t-test) and BI-1 for days 2, 4, and 7. BI-1 was not detected in the stool on day 9 post-infection (*).

Next, we performed a powered animal study comparing the virulence of the LT-027 isolate GV148 (N = 8) to BI-1 (N = 6). All GV148-infected animals displayed similar disease progression and succumbed to disease between 24-40-hours post-infection (average time to death was 32 hours; Figure 4C). BI-1-infected animals showed a trend towards earlier death (24-30 hours), but this was not statistically significant relative to the mortality kinetics for GV148-infected animals. Control hamsters infected with the non-toxigenic strain T7, expectedly, survived the infection. These data suggest that, despite producing significantly lower amounts of toxin, LT-027 strains exhibit robust virulence in the hamster model.

### LT-027 *C. difficile* are shed in greater numbers and persist longer than strain BI-1 in infected mice

In the hamster model, *C. difficile-*infected animals succumb to infection within 48 hours, potentially masking strain-specific differences in disease progression. We therefore used a non-lethal mouse model to assess colonization differences between LT-027 strains and BI-1. Antibiotic-treated C57BL/6 mice were orally administered either PBS, or spores of strains BI-1, or an LT-027 strain (GV106, GV135, or GV148). *C. difficile* was detected in the stools of all animals by day 2 post-infection, with a 100-fold higher initial burden in mice infected with BI-1 compared to those treated with LT-027 (Figure 4D). Although stool *C. difficile* burdens increased in all animals by day 4 post-infection, we observed significantly higher pathogen counts for all LT-027-infected mice relative to BI-1. Subsequently, BI-1 burdens in the stool decreased by day 7, with complete clearance by day 9 post-infection. In sharp contrast, LT-027 strains persisted at high levels in the stool on days 7 and 9 post-infection. These results suggest that LT-027 strains colonize better, and persist longer, than BI-1.

### *C. difficile* LT-027 strains express unique proteomes

The increased stool burdens for GV106, GV135, or GV148, relative to BI-1 suggested that LT-027 strains may have unique characteristics, beyond decreased toxin levels, that facilitate colonization. To identify contributing factors, we performed a comparative, quantitative, mass spectrometry-based proteomics analysis on 3 LT-027 isolates (GV106, GV135, and GV148) relative to BI-1. Despite belonging to the same lineage as BI-1 (RT027), there were marked protein abundance differences between each of the LT-027 isolates and BI-1 at mid-logarithmic growth phase (Supplemental Figure S2, Supplemental Table S5-S7). In GV106, one protein, a Sigma 54 interacting transcription anti-terminator, was significantly upregulated, and 63 proteins were downregulated (Table S6). GV135 had 5 proteins significantly upregulated, and 5 proteins downregulated (Table S5). GV148 had 8 proteins significantly upregulated, and 6 proteins downregulated (Table S4).

Importantly, and collectively, there were notable alterations in the abundances of cell wall-associated proteins in the LT-027 strains. GV135 and GV148 had significantly increased abundance of D-alanyl-D-serine ligase (CDBI-1_08005) and D-alanyl-D-alanine carboxypeptidase (CDBI-1_08010) (Supplementary Table 4 and 5). GV106 D-alanyl-D-serine ligase and D-alanyl-D-alanine carboxypeptidase were downregulated compared to BI-1, though not significantly so (Supplementary Table 6). Additionally, all three strains had decreased abundance of D-alanyl-D-alanine ligase (CDBI-1_06520) and, in the case of GV106, SlpA was downregulated. Further, the S-layer precursor protease, Cwp84 (CDBI-1_13585), was less abundant in the LT-027 than in BI-1.

### *C. difficile* LT-027 strain express unique surface antigens

The marked alteration in abundance of select LT-027 proteins, coupled with increased mouse gut colonization, suggested possible differences in bacterial cell surface architecture and, consequently, antigen profile, relative to BI-1. To assess this, membrane-bound protein fractions of LT-027 and comparator strains were extracted and probed with serum harvested from infected mice. Serum from LT-027-infected animals recognized unique cell-surface proteins (∼75 kDa) from LT-027 membrane fractions (yellow arrow), but not from the high-toxin producing RT027 (BI-1) membrane fraction (Figure 5). This difference, however, was not recapitulated when probing membrane fractions of high-toxin (GV599) and low-toxin (GV1002) isolates from an unrelated *C. difficile* lineage (RT106; Figure 5A and 5B), indicating potential ribotype-specificity of cell-surface architecture.

**FIGURE 5:**
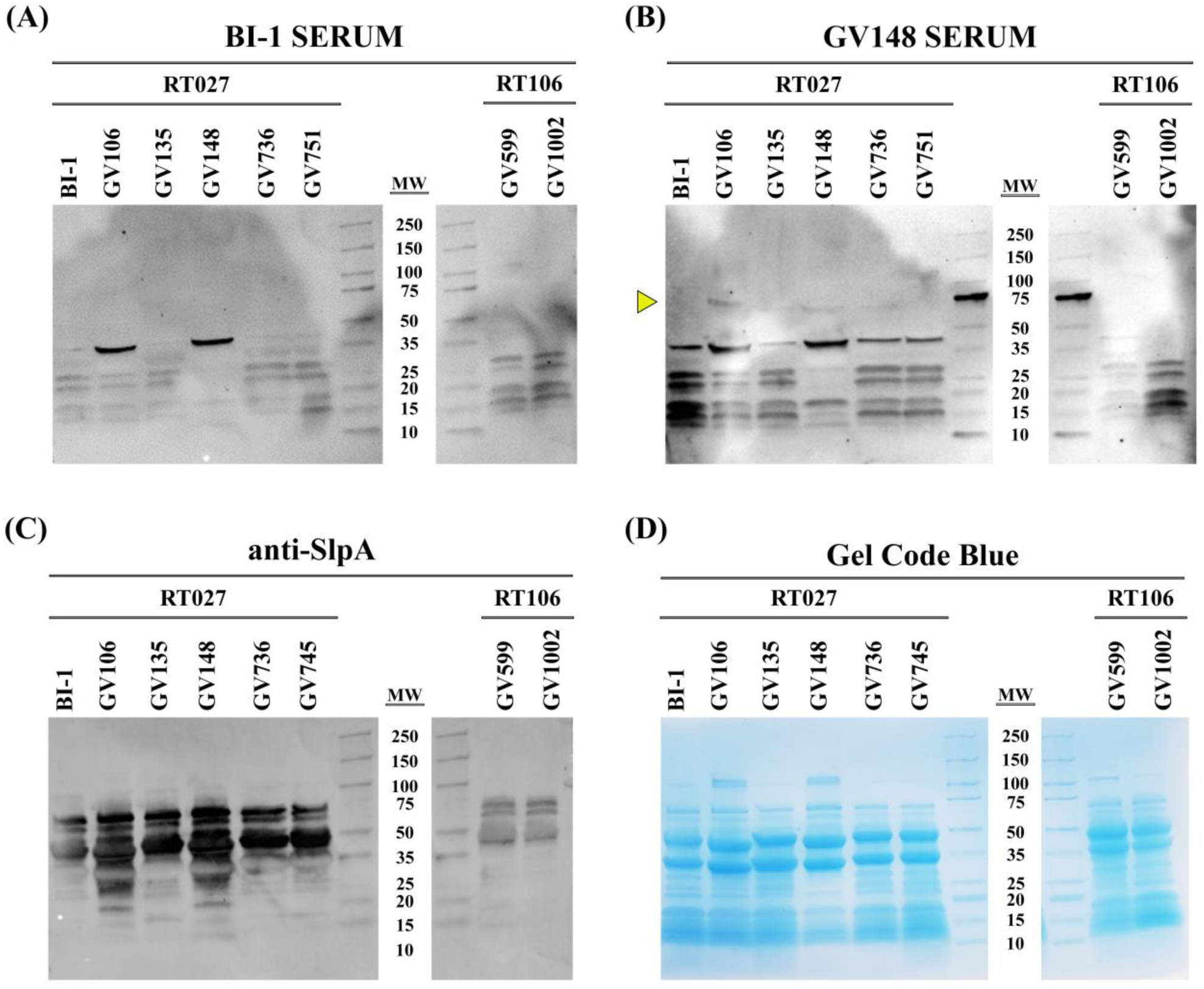
LT RT027strains exhibit a unique antigen profile. PAGE-separated crude *C. difficile* surface layer extracts (30 µg/lane) from high- (BI-1 and GV751) and low-toxin strains (BV148, GV736, GV745) were probed with serum isolated from mice infected with the high-toxin RT027 strain BI-1 (A), LT-027 strain GV148 (B), or anti-SlpA antibodies (C). Protein quantities shown with Gel Code Blue stain (D). GV148 antiserum detects a unique band (yellow arrow, ∼75 kDa) in surface extracts of LT-027, but not the high-toxin strain BI-1. Further, this band is not detected when probed with BI-1-infected mouse serum.

### LT-027 *C. difficile* share unique genomic content not found in the reference RT027

To determine if there is genetic content unique to the low-toxin strains, we compared whole genomes of 17 LT-027 strains to the reference RT027 strain BI-1; all isolates had similar genome sizes (4.1 – 4.28MB). LT-027 strains harbored no obvious unique DNA islands, or toxin/antibiotic resistance genes. All LT-027 strains harbored binary toxin. In all sequenced LT-027 strains, the PaLoc was 100% conserved relative to BI-1, including the 18bp deletion in *tcdC*. Known regulators of toxin gene expression (TcdR, CodY, CcpA, SigD, and RstA [32]) were also 100% conserved among all RT027 strains.

Next, an alignment-free composite vector tree was constructed (CVTree 3.0) from 15 sequenced LT-027 strains, 3 GV-RT027 strains, and 10 publicly available RT027 strains. We observed that LT-027 strains (red) clade separately from reference/high-toxin strains (black) (Supplemental Figure S1). This suggests there are common genetic elements shared between LT-027 strains. To determine this, we generated a pan-genome for LT-027 strains (N = 10) and compared this to the pan-genome of high-toxin RT027 strains (N = 10), including R20291, CD196, BI-1, and BI-8. This analysis revealed 66 genes unique to the low-toxin pan-genome (Supplemental Table S4). Of these, 10 genes were predicted to encode cell surface-associated molecules (Table 1). It is possible that one, or several, of these genes are responsible for the increased colonization seen in the murine model.

**Table 1:**
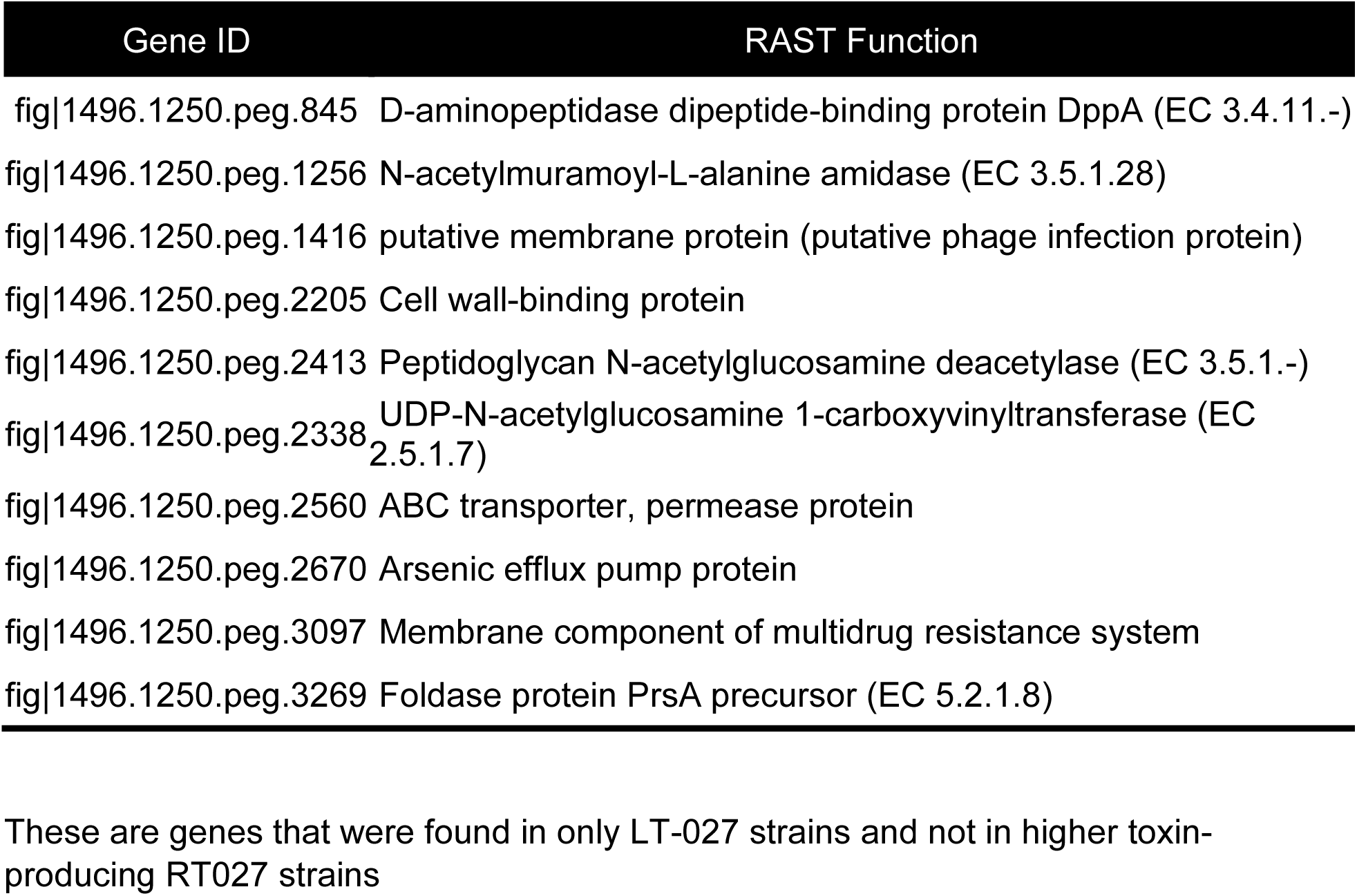
Cell-wall-associated genes unique to LT-027 strains. These are genes that were found in only LT-027 strains and not in higher toxin-producing RT027 strains

The studies above describe distinct differences of low-toxin strains in within a single *C. difficile* lineage (RT027), raising the possibility that the observed phenotypes and genotypes are ribotype-specific. Therefore, we assessed high- and low-toxin-producing strains of another USA-dominant *C. difficile* lineage, RT106. Genetic differences between low-toxin RT106 (LT-106) and high-toxin RT106 strains were not as stark as their RT027 counterparts, with fewer genes (n=18) identified in the LT-106 pangenome (Supplemental Table S8). Only one gene (predicted to encode an RNA-binding protein) was common between the LT-027 and LT-106 pangenomes. Unlike the LT-027 strains, LT-106 strains were interspersed between high-toxin RT106 strains in a composite vector tree (Supplemental Figure S4). Despite these differences, and similar to LT-027, the LT-106 strains also exhibited increased colonization in mice (Supplemental Figure S4). Furthermore, immunoblots using serum of LT-106 infected animals revealed unique bands in LT-106 membrane fractions but not in high-toxin RT106, or any RT027, membrane fractions (Supplemental Figure S5).

## DISCUSSION

Collectively, our data suggest that low-toxin strains isolated from discrepant patient stool samples consistently express >10-fold lower toxin levels than reference high-toxin strains, irrespective of ribotype. LT-027 strains, specifically, appear to clade separately from other RT027 strains. Low-toxin strains have altered cell surfaces, though these differences are not consistent across ribotypes, resulting in unique serum reactivity. Interestingly, these low-toxin strains are able to colonize and persist in mouse intestines and exhibit robust lethality in the hamster model of infection.

While it is well established that the glucosylating toxins TcdA and TcdB are essential for *C. difficile* virulence [13,20,33-36], there is no evidence to suggest that a threshold toxin amount is required to elicit diarrheagenic CDI symptoms. When first described the increased virulence of epidemic-associated RT027 strains was attributed to an increase in toxin production [37]. Multiple studies, however, found no evidence for consistent production of higher TcdA and TcdB amounts by RT027 strains compared to other ribotypes [31,38-42]. Our studies also confirm that clinical isolates, including RT027 strains, vary significantly in the production of TcdA and TcdB.

In the clinical setting, numerous studies have reported correlations between stool toxin levels and disease severity [43,44]. These observations underpin the recommendation that a diagnosis of CDI should be based on positive detection of TcdA/TcdB in the stool via EIA, the limited sensitivity of this assay serving as a de facto diagnostic threshold [41,45]. The rationale is that this would exclude instances of non-CDI diarrhea, but with happenstance *C. difficile* colonization. In our studies, however, fully toxigenic *C. difficile* strains were recovered from >30% of toxin EIA-negative stool specimens, all from symptomatic patients. Similarly, Erb et al noted that 42.9% (N = 206) of patient samples were CDI-positive via toxin culture despite failing the stool EIA test for toxin [41]. Of particular concern is that they did not find any difference in mortality or CDI recurrence between EIA Tox^-^ and EIA Tox^+^ groups. Other studies have also reported similar findings and have additionally noted toxin EIA discrepancy for multiple *C. difficile* ribotypes [38,45,46].

Considering the spectrum of host susceptibility, and of *C. difficile* strain variability, it is challenging to demarcate the window within which attributable disease manifests (‘pathogenicity’) [47]. The Damage-Response Framework, proposed by Pirofski and Casadevall [48] highlights the role of the host, and may be useful in understanding the consequences of *C. difficile* infection. While some patients may have CDI despite undetectable stool toxin levels, others harboring toxigenic *C. difficile* with detectable stool toxin may be asymptomatic. Our studies demonstrating the robust virulence of low-toxin-producing RT027 strains highlight, rather than resolve, this dilemma.

From a selection standpoint, it is possible that low-toxin *C. difficile* strains may proliferate or become endemic in hospital settings (especially where there is reliance on stool toxin tests) precisely because these strains escape attention or treatment. Toxin expression is highly regulated and intricately linked to *C. difficile* metabolism [49-54] and the metabolic cost of producing TcdA and TcdB is high. There are no data to suggest a benefit to *C. difficile* producing high quantities of these large molecules. In their study of clinical *Staphylococcus aureus* strains, Laabei et al demonstrated that low-toxicity isolates had a higher propensity to cause bacteremia than highly toxic strains [55]. Their work suggested that the high energy requirements of toxin production might exact a fitness cost on high-toxin isolates within the host. Interestingly, Hollands et al noted that a genetic switch to hypervirulence in Group A *Streptococcus* strains decreased the colonization capacity of these strains [56]. These findings are consistent with the idea that bacterial virulence is a trade-off between toxicity, relative fitness, and transmissibility.

Our studies suggest that low-toxin strains may harbor hyper-colonization potential compared to high toxin strains. Despite being the same ribotype, low-toxin strains harbor unique genes and have an altered proteome relative to high-toxin strains. Specifically, LT-027 strains elaborate a distinct surface architecture that elicits an equally-distinct serum reactivity which may contribute to the robust gut colonization and persistence we observed. This difference, however, does not seem to involve peptidoglycan remodeling since vancomycin susceptibility is statistically similar between the RT027 strains we tested. Additionally, the pangenomes of LT-027 and those of unrelated *C. difficile* lineages like RT106, share only one gene. This lack of inter-lineage gene-content overlap is consistent with the enormous plasticity of the *C. difficile* genome [57], and may suggest convergent evolutionary/genetic mechanisms that, on the whole, potentiate stable low-toxin and colonization phenotypes.

Given the prevalence of low-toxin strains in clinical settings, both in our study and in other reports [11,38,58], failure to detect toxin in the stool of *C. difficile*-positive patients should be interpreted with caution. In their study, Reigadas et al reported 12.7% of CDI cases (n = 26 of 204 total) would have been missed due to a lack of clinical suspicion despite these patients exhibiting mild-to-moderate symptoms [59]. These 26 patients experienced similar duration of diarrhea as patients with suspected CDI cases, but only 30.8% received CDI treatment and 11.5% had recurrent infections.

Taken together, work by us and others indicate that CDI cases are likely being missed and underreported to an unknown degree. Though dogma suggests that these cases result in mild symptoms, our work demonstrates the ability of low-toxin strains to potentiate fulminant disease. Since these strains were isolated from symptomatic, diarrheal patients, it is imperative that discrepant *C. difficile* strains are not overlooked.

## ACKNOWLEDGEMENTS

This work was supported by funding from the National Institutes of Health [R33AI121590531(GV)], the US Department of Veterans Affairs [IK6BX003789(GV Research Career Scientist Award); I01BX001183(GV Merit Award)] and a University of Arizona Research, Innovation & Impact Award [UA5833256(GV)]. Ribotyping work was performed with the support of the University of Arizona Genetics Core, University of Arizona, Tucson, AZ.

