## Supplementary Data for "Low-Toxin *Clostridioides difficile* RT027 Strains Exhibit Robust Virulence"

[1]School of Animal and Comparative Biomedical Sciences, The University of Arizona, Tucson, AZ, USA, [2]Department of Pediatrics, The University of Arizona College of Medicine, Tucson, AZ, USA, [3]Mayo Clinic, Phoenix, AZ, [4]Department of Immunobiology, The University of Arizona College of Medicine, Tucson, AZ, USA, [5]BIO5 Institute for Collaborative Research, The University of Arizona, Tucson, AZ, [6]Southern Arizona VA Healthcare System, Tucson, AZ[4].

*Correspondence to:

Gayatri Vedantam, Ph.D.
University of Arizona
School of Animal & Comparative Biomedical Sciences
1117, E. Lowell St. Bldg. 90, Room
Tucson, AZ 85721


**Supplemental Table S1: Primers Used In this Study**

| Primer name | Gene Target | Sequence | Ref |
|---|---|---|---|
| RIBO-F | 16S rDNA | 5′-GCTGGATCACCTCCTTTCTAAG | Janežič et al. *J Clin Microbiol* (2011) |
| RIBO-R | 23S rDNA | 5′-TGACCAGTTAAAAAGGTTTGATAGATT | |
| B1C | *tcdB* | 5' - AGAAAATTTTATGAGTTTAGTTAATAGAAA | Rupnik et al. *FEMS Microbiol Lett* (2006) |
| B2N | *tcdB* | 5' - CAGATAATGTAGGAAGTAAGTCTATAG | |
| A1C | *tcdA* | 5' - GGAGGTTTTTATGTCTTTAATATCTAAAGA | |
| A2N | *tcdA* | 5' - CCCTCTGTTATTGAGGTAGTACATTTA | |
| cdtA-F | *cdtA* | 5′-TGAACCTGGAAAAGGTGATG | Pituch et al. *J Med Microbiol* (2005) |
| cdtA-R | *cdtA* | 5′-AGGATTATTTACTGGACCATTTG | |
| cdtB-F | *cdtB* | 5′-CTTAATGCAAGTAAATACTGAG | |
| cdtB-R | *cdtB* | 5′-AACGGATCTCTTGCTTCAGTC | |
| Tim2 | *tcdC* | 5’-GCACCTCATCACCATCTTCA | Cohen et al. *J Infect Dis* (2000) |
| Struppi2 | *tcdC* | 5’-TGAAGACCATGAGGAGGTCAT | |

**Supplemental Table S2: GenBank Accession Numbers of LT-027 Isolates**

| Strain | NCBI Accession Numbers | | Coverage |
|---|---|---|---|
| | Biosample | BioProject | |
| GV106 | SAMN26752631 | PRJNA817265 | 630x |
| GV135 | SAMN26752621 | PRJNA817265 | 2060x |
| GV144 | SAMN26752619 | PRJNA817265 | 65x |
| GV145 | SAMN26752625 | PRJNA817265 | 142x |
| GV147 | SAMN26752628 | PRJNA817265 | 69x |
| GV148 | SAMN26752630 | PRJNA817265 | 751x |
| GV153 | SAMN26752620 | PRJNA817265 | 51x |
| GV155 | SAMN26752626 | PRJNA817265 | 142x |
| GV161 | SAMN26752624 | PRJNA817265 | 134x |
| GV163 | SAMN26752633 | PRJNA817265 | 145x |
| GV165 | SAMN26752635 | PRJNA817265 | 157x |
| GV175 | SAMN26752627 | PRJNA817265 | 120x |
| GV178 | SAMN26752623 | PRJNA817265 | 118x |
| GV183 | SAMN26752629 | PRJNA817265 | 98x |
| GV736 | SAMN26752622 | PRJNA817265 | 195x |
| GV745 | SAMN26752634 | PRJNA817265 | 120x |
| GV752 | SAMN26752632 | PRJNA817265 | 318x |

### Supplemental Table S3: MIC of Select Antibiotics of High- and Low-Toxin Strains

| MIC (µg/mL) | Cefotaxime | Rifampicin | Levofloxacin | Metronidazole | Vancomycin |
|---|---|---|---|---|---|
| **BI-1** | >16 µg/mL* | 1 µg/mL | >32 µg/mL* | >256 µg/mL* | 3 µg/mL |
| **GV106** | >16 µg/mL* | 1 µg/mL | >32 µg/mL* | >256 µg/mL* | 3 µg/mL |
| **GV135** | 0.75 µg/mL* | 3 µg/mL | 0.75 µg/mL | >256 µg/mL* | 3 µg/mL |
| **GV148** | >16 µg/mL | >32 µg/mL* | >32 µg/mL* | 1 µg/mL | 4 µg/mL |

Units are in µg/mL.
*Denotes that the strain is highly resistant.

### Supplemental Table S4: Unique genes harbored by LT-027 strains

| Gene ID | Function |
|---|---|
| fig\|1496.1250.peg.42 | FIG002813: LPPG:FO 2-phospho-L-lactate transferase like, CofD-like |
| fig\|1496.1250.peg.146 | FIG00514768: hypothetical protein |
| fig\|1496.1250.peg.155 | Anaerobic sulfite reductase subunit B |
| fig\|1496.1250.peg.164 | N-acetylmannosamine-6-phosphate 2-epimerase (EC 5.1.3.9) |
| fig\|1496.1250.peg.173 | ATPase associated with various cellular activities, AAA_5 |
| fig\|1496.1250.peg.198 | FIG00522519: hypothetical protein |
| fig\|1496.1250.peg.371 | tRNA and rRNA cytosine-C5-methylases |
| fig\|1496.1250.peg.394 | FIG00520447: hypothetical protein |
| fig\|1496.1250.peg.407 | FIG00512777: hypothetical protein |
| fig\|1496.1250.peg.408 | FIG00515422: hypothetical protein |
| fig\|1496.1250.peg.470 | Cytosol nonspecific dipeptidase (EC 3.4.13.18) |
| fig\|1496.1250.peg.661 | Osmosensitive K+ channel histidine kinase KdpD (EC 2.7.3.-) |
| fig\|1496.1250.peg.755 | Chloride channel protein |
| fig\|1496.1250.peg.845 | D-aminopeptidase dipeptide-binding protein DppA (EC 3.4.11.-) |
| fig\|1496.1250.peg.965 | LSU ribosomal protein L13p (L13Ae) |
| fig\|1496.1250.peg.1001 | DNA topoisomerase I (EC 5.99.1.2) |
| fig\|1496.1250.peg.1050 | Translocation-enhancing protein TepA |
| fig\|1496.1250.peg.1145 | RNA polymerase sporulation specific sigma factor SigE |
| fig\|1496.1250.peg.1158 | Acyl-CoA:1-acyl-sn-glycerol-3-phosphate acyltransferase (EC 2.3.1.51) |
| fig\|1496.1250.peg.1173 | 3-dehydroquinate synthase (EC 4.2.3.4) |
| fig\|1496.1250.peg.1221 | AzlC family protein |
| fig\|1496.1250.peg.1256 | N-acetylmuramoyl-L-alanine amidase (EC 3.5.1.28) |
| fig\|1496.1250.peg.1324 | Transcriptional regulator |
| fig\|1496.1250.peg.1335 | Hydantoinase/oxoprolinase family protein |
| fig\|1496.1250.peg.1416 | putative membrane protein (putative phage infection protein) |
| fig\|1496.1250.peg.1419 | Rubrerythrin |
| fig\|1496.1250.peg.1455 | FIG00519734: hypothetical protein |
| fig\|1496.1250.peg.1483 | FIG00514478: hypothetical protein |
| fig\|1496.1250.peg.1580 | Peptide methionine sulfoxide reductase MsrA (EC 1.8.4.11) / Peptide methionine sulfoxide reductase MsrB (EC 1.8.4.12) |
| fig\|1496.1250.peg.1736 | V-type ATP synthase subunit D (EC 3.6.3.14) |
| fig\|1496.1250.peg.2084 | Stage III sporulation protein AB |
| fig\|1496.1250.peg.2198 | SSU ribosomal protein S20p |
| fig\|1496.1250.peg.2205 | Cell wall-binding protein |
| fig\|1496.1250.peg.2206 | Uracil-DNA glycosylase, family 1 |
| fig\|1496.1250.peg.2337 | FIG013354: hypothetical protein |
| fig\|1496.1250.peg.2338 | UDP-N-acetylglucosamine 1-carboxyvinyltransferase (EC 2.5.1.7) |
| fig\|1496.1250.peg.2342 | MreB-like protein (Mbl protein) |
| fig\|1496.1250.peg.2378 | hypothetical protein |
| fig\|1496.1250.peg.2379 | RNA polymerase sporulation specific sigma factor SigK |
| fig\|1496.1250.peg.2413 | Peptidoglycan N-acetylglucosamine deacetylase (EC 3.5.1.-) |
| fig\|1496.1250.peg.2440 | ATP phosphoribosyltransferase (EC 2.4.2.17) |
| fig\|1496.1250.peg.2466 | FIG00522895: hypothetical protein |
| fig\|1496.1250.peg.2476 | FIG006036: phage encoded DNA polymerase I (EC 2.7.7.7) |
| fig\|1496.1250.peg.2527 | HNH homing endonuclease |

| | |
|---|---|
| fig\|1496.1250.peg.2560 | ABC transporter, permease protein |
| fig\|1496.1250.peg.2609 | Formate dehydrogenase H (EC 1.2.1.2) @ selenocysteine-containing |
| fig\|1496.1250.peg.2611 | Response regulator |
| fig\|1496.1250.peg.2670 | Arsenic efflux pump protein |
| fig\|1496.1250.peg.2790 | Dihydropyrimidinase (EC 3.5.2.2) @ D-hydantoinase (EC 3.5.2.2) |
| fig\|1496.1250.peg.2815 | Transcriptional regulator, MerR family |
| fig\|1496.1250.peg.2862 | Transcriptional regulator, MecI family |
| fig\|1496.1250.peg.2904 | FIG004454: RNA binding protein |
| fig\|1496.1250.peg.2992 | FIG00516716: hypothetical protein |
| fig\|1496.1250.peg.3048 | Sarcosine reductase component B beta subunit (EC 1.21.4.3) |
| fig\|1496.1250.peg.3092 | Hydrolase, HAD superfamily |
| fig\|1496.1250.peg.3097 | Membrane component of multidrug resistance system |
| fig\|1496.1250.peg.3102 | Beta-glucoside bgl operon antiterminator, BglG family |
| fig\|1496.1250.peg.3114 | RecA regulator RecX |
| fig\|1496.1250.peg.3123 | Phosphate regulon transcriptional regulatory protein PhoB (SphR) |
| fig\|1496.1250.peg.3218 | FIG00518106: hypothetical protein |
| fig\|1496.1250.peg.3269 | Foldase protein PrsA precursor (EC 5.2.1.8) |
| fig\|1496.1250.peg.3342 | Glycerol-3-phosphate regulon repressor GlpR |
| fig\|1496.1250.peg.3462 | ATP-dependent DNA helicase UvrD/PcrA |
| fig\|1496.1250.peg.3656 | FIG001621: Zinc protease |
| fig\|1496.1250.peg.3859 | putative PTS system, IIa component |
| fig\|1496.1250.peg.3966 | Ribonucleotide reductase of class III (anaerobic), activating protein (EC 1.97.1.4) |

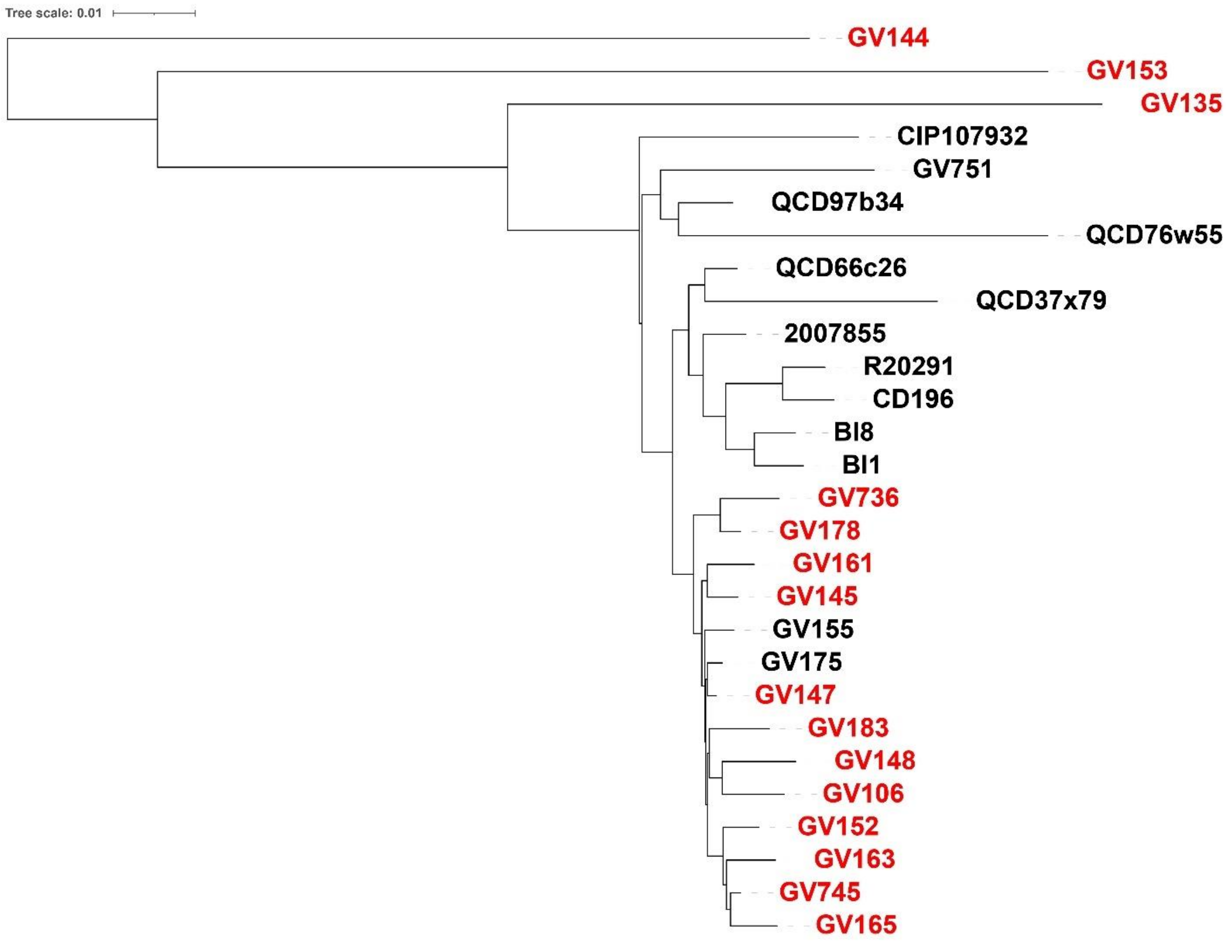


**<u>Supplemental Figure S1: Phylogenetic Comparison of High- and Low-Toxin RT027 Strains</u>**

Whole genome comparison was done between clinically derived, low-toxin producing (n = 15; red) and high toxin producing strains (n = 13; black) using CVTree3 (http://tlife.fudan.edu.cn/cvtree3/). The high toxin strains (black) were previously published (except for GV155 and GV175).

Whole genome comparison of these 28 strains show distinct clades that separate the high and low toxin strains. Furthermore, there are, at minimum, 80 SNP differences between the most closely related low toxin strains indicating that these clinical strains are not clonal.

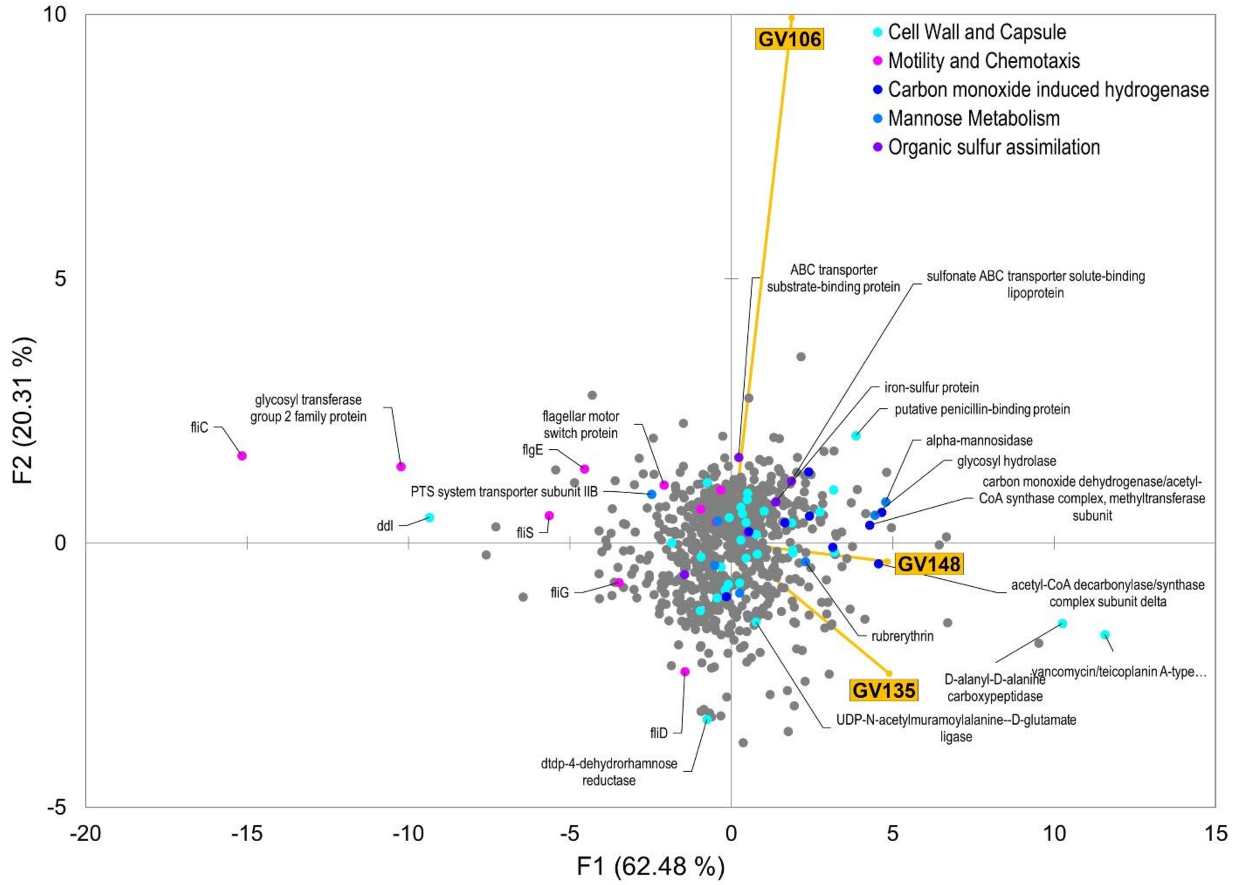


**Supplemental Figure S2: PCA of LT-027 protein abundances relative to BI-1.**

Protein abundances of low-toxin strains, at mid-log growth, were compared to BI-1. Each dot represents a protein hit; colored dots are proteins involved in selected biological processes and were statistically significant differences in abundance. Several proteins that were differentially abundant belonged to cell wall and capsule biology.

**Supplemental Table S5: Statistically significant alterations in protein abundances on GV148 compared to BI-1**

| Locus | Locus ID | Strain | Name | GV148:BI-1 | |
|---|---|---|---|---|---|
| | | | | Fold Change (Log2 Ratio) | P-value (-Log10) |
| CDBI1_08005 | CDBI1_08005 | Peptoclostridium difficile BI1 | vancomycin/teicoplanin A-type resistance protein [D-alanine-D-serine Ligase] | ***2.950*** | ***4.074*** |
| CDBI1_11295 | CDBI1_11295 | Peptoclostridium difficile BI1 | NAD-dependent 4-hydroxybutyrate dehydrogenase | ***2.432*** | ***1.452*** |
| CDBI1_08010 | CDBI1_08010 | Peptoclostridium difficile BI1 | D-alanyl-D-alanine carboxypeptidase | ***2.285*** | ***3.328*** |
| CDBI1_02690 | CDBI1_02690 | Peptoclostridium difficile BI1 | NADP-dependent glyceraldehyde-3-phosphate dehydrogenase | ***2.166*** | ***2.014*** |
| CDBI1_13615 | CDBI1_13615 | Peptoclostridium difficile BI1 | S-layer precursor protein [Surface layer protein A] | ***1.847*** | ***6.220*** |
| obgE | CDBI1_05225 | Peptoclostridium difficile BI1 | GTPase CgtA | ***1.355*** | ***2.353*** |
| CDBI1_07840 | CDBI1_07840 | Peptoclostridium difficile BI1 | putative O-acetylserine sulfhydrylase | ***1.076*** | ***2.863*** |
| CDBI1_13635 | CDBI1_13635 | Peptoclostridium difficile BI1 | hypothetical protein CDBI1_13635 [Calcium-binding adhesion protein] | ***1.010*** | ***1.675*** |
| CDBI1_01045 | CDBI1_01045 | Peptoclostridium difficile BI1 | chaperonin GroEL | 0.718 | ***1.517*** |
| CDBI1_05845 | CDBI1_05845 | Peptoclostridium difficile BI1 | cysteine desulfurase | 0.571 | ***1.398*** |
| CDBI1_11245 | CDBI1_11245 | Peptoclostridium difficile BI1 | sigma-54 interacting transcription antiterminator | 0.425 | ***1.398*** |
| CDBI1_04765 | CDBI1_04765 | Peptoclostridium difficile BI1 | butyryl-CoA dehydrogenase | 0.345 | ***1.333*** |
| CDBI1_14240 | CDBI1_14240 | Peptoclostridium difficile BI1 | bifunctional acetaldehyde-CoA/alcohol dehydrogenase | 0.093 | ***7.116*** |
| CDBI1_00265 | CDBI1_00265 | Peptoclostridium difficile BI1 | prolyl-tRNA synthetase | -0.332 | ***1.595*** |
| thrS | CDBI1_02630 | Peptoclostridium difficile BI1 | threonyl-tRNA synthetase | -0.452 | ***1.564*** |
| CDBI1_10630 | CDBI1_10630 | Peptoclostridium difficile BI1 | cyclomaltodextrinase | -0.465 | ***1.786*** |
| CDBI1_11470 | CDBI1_11470 | Peptoclostridium difficile BI1 | quinolinate synthetase | -0.651 | ***1.705*** |

| | | | | | |
|---|---|---|---|---|---|
| rplB | CDBI1_00425 | Peptoclostridium difficile BI1 | 50S ribosomal protein L2 | -0.731 | ***1.678*** |
| CDBI1_17325 | CDBI1_17325 | Peptoclostridium difficile BI1 | methionyl-tRNA synthetase | ***-1.023*** | ***1.520*** |
| prdB | CDR20291_3101 | Peptoclostridium difficile R20291 | proline reductase | ***-1.050*** | ***1.560*** |
| CDBI1_03635 | CDBI1_03635 | Peptoclostridium difficile BI1 | formate acetyltransferase | ***-1.116*** | ***6.190*** |
| guaA | CDBI1_01065 | Peptoclostridium difficile BI1 | GMP synthase | ***-1.342*** | ***1.795*** |
| CDBI1_04210 | CDBI1_04210 | Peptoclostridium difficile BI1 | ABC transporter substrate-binding protein | ***-3.681*** | ***2.001*** |
| CDBI1_13585 | CDBI1_13585 | Peptoclostridium difficile BI1 | cell surface-associated cysteine protease [Cwp84] | ***-3.681*** | ***2.001*** |

Red indicates high abundance; green indicates less abundance. Increasing fold changes are reported as brighter color.
Fold changes reported in Log2 values.
P-values reported as -log10 values.

**Supplemental Table S6: Statistically significant alterations in protein abundances of GV135 compared to BI-1.**

| Locus | Locus ID | Strain | Name | GV135:BI-1 | |
|---|---|---|---|---|---|
| | | | | Fold Change (Log2 Ratio) | P-value (-Log10) |
| CDBI1_08005 | CDBI1_08005 | Peptoclostridium difficile BI1 | vancomycin/teicoplanin A-type resistance protein [D-alanyl-D-serine ligase] | ***3.030*** | ***2.495*** |
| CDBI1_08010 | CDBI1_08010 | Peptoclostridium difficile BI1 | D-alanyl-D-alanine carboxypeptidase | ***2.339*** | ***4.292*** |
| CDBI1_11295 | CDBI1_11295 | Peptoclostridium difficile BI1 | NAD-dependent 4-hydroxybutyrate dehydrogenase | ***2.033*** | ***1.437*** |
| obgE | CDBI1_05225 | Peptoclostridium difficile BI1 | GTPase CgtA | ***1.595*** | ***2.361*** |
| CDBI1_01045 | CDBI1_01045 | Peptoclostridium difficile BI1 | chaperonin GroEL | ***1.143*** | ***2.314*** |
| CDBI1_07840 | CDBI1_07840 | Peptoclostridium difficile BI1 | putative O-acetylserine sulfhydrylase | **0.704** | ***2.829*** |
| CDBI1_05845 | CDBI1_05845 | Peptoclostridium difficile BI1 | cysteine desulfurase | **0.611** | ***1.819*** |
| cysS | CDBI1_00275 | Peptoclostridium difficile BI1 | cysteinyl-tRNA synthetase | **0.611** | ***1.333*** |
| CDBI1_13280 | CDBI1_13280 | Peptoclostridium difficile BI1 | putative serine hydroxymethyltransferase | **0.558** | ***1.429*** |
| CDBI1_13425 | CDBI1_13425 | Peptoclostridium difficile BI1 | phosphoenolpyruvate-protein phosphotransferase | **0.505** | ***2.426*** |
| dnaK | CDBI1_11965 | Peptoclostridium difficile BI1 | molecular chaperone DnaK | **0.306** | ***1.766*** |
| CDBI1_14240 | CDBI1_14240 | Peptoclostridium difficile BI1 | bifunctional acetaldehyde-CoA/alcohol dehydrogenase | **0.120** | ***4.820*** |
| CDBI1_15490 | CDBI1_15490 | Peptoclostridium difficile BI1 | phosphoglyceromutase | **-0.532** | ***1.320*** |
| CDBI1_09460 | CDBI1_09460 | Peptoclostridium difficile BI1 | hypothetical protein CDBI1_09460 | **-0.558** | ***1.840*** |
| CDBI1_03470 | CDBI1_03470 | Peptoclostridium difficile BI1 | acetyl-CoA decarbonylase/synthase complex subunit gamma | **-0.651** | ***1.684*** |
| CDBI1_03635 | CDBI1_03635 | Peptoclostridium difficile BI1 | formate acetyltransferase | **-0.704** | ***1.407*** |
| prfA | CDBI1_17040 | Peptoclostridium difficile BI1 | peptide chain release factor 1 | ***-0.917*** | ***1.308*** |

| | | | | | |
|---|---|---|---|---|---|
| CDBI1_15485 | CDBI1_15485 | Peptoclostridium difficile BI1 | enolase | *-1.621* | ***2.186*** |
| CDBI1_13595 | CDBI1_13595 | Peptoclostridium difficile BI1 | cell surface protein [Cell wall binding repeat-containing protein] | ***-2.684*** | ***1.858*** |
| CDBI1_04210 | CDBI1_04210 | Peptoclostridium difficile BI1 | ABC transporter substrate-binding protein | ***-4.292*** | ***2.180*** |
| CDBI1_13585 | CDBI1_13585 | Peptoclostridium difficile BI1 | cell surface-associated cysteine protease [Cell wall binding protein Cwp84] | ***-4.292*** | ***2.180*** |

Red indicates high abundance; green indicates less abundance. Increasing fold changes are reported as brighter color.
Fold changes reported in Log2 values.
P-values reported as -log10 values.

**Supplemental Table S7: Statistically significant alterations in protein abundances of GV106 compared to BI-1.**

| Locus | Locus ID | Strain | Name | GV106:BI-1 Fold Change (Log2 Ratio) | GV106:BI-1 P-value (-Log10) |
|---|---|---|---|---|---|
| CDBI1_11245 | CDBI1_11245 | Peptoclostridium difficile BI1 | sigma-54 interacting transcription antiterminator | *0.997* | ***1.549*** |
| addA | QO7_0989 | Peptoclostridium difficile F314 | helicase-exonuclease AddAB, AddA subunit | 0.385 | ***2.995*** |
| CDBI1_15555 | CDBI1_15555 | Peptoclostridium difficile BI1 | diaminopropionate ammonia-lyase | 0.319 | ***2.891*** |
| CDBI1_09370 | CDBI1_09370 | Peptoclostridium difficile BI1 | acetyl-coenzyme A carboxylase carboxyl transferase subunit alpha | -0.080 | ***1.495*** |
| CDBI1_12720 | CDBI1_12720 | Peptoclostridium difficile BI1 | cell-division initiation protein | -0.146 | ***1.480*** |
| CDBI1_16555 | CDBI1_16555 | Peptoclostridium difficile BI1 | HPr kinase/phosphorylase | -0.372 | ***2.049*** |
| CDBI1_01350 | CDBI1_01350 | Peptoclostridium difficile BI1 | flagellum-specific ATP synthase | -0.598 | ***2.116*** |
| CDBI1_17110 | CDBI1_17110 | Peptoclostridium difficile BI1 | bifunctional tetrapyrrole (Corrin/Porphyrin) methylase/nucleoside triphosphate pyrophosphohydrolase | -0.625 | ***1.538*** |
| CDBI1_13605 | CDBI1_13605 | Peptoclostridium difficile BI1 | S-layer protein | -0.691 | ***1.679*** |
| CDBI1_11885 | CDBI1_11885 | Peptoclostridium difficile BI1 | tRNA binding protein | *-0.877* | ***1.483*** |
| CDBI1_15775 | CDBI1_15775 | Peptoclostridium difficile BI1 | amidohydrolase | *-1.143* | ***2.345*** |
| CDBI1_17610 | CDBI1_17610 | Peptoclostridium difficile BI1 | hypothetical protein CDBI1_17610 | *-1.342* | ***1.918*** |
| rpsE | CDBI1_00495 | Peptoclostridium difficile BI1 | 30S ribosomal protein S5 | *-1.541* | ***1.718*** |
| CDBI1_12555 | CDBI1_12555 | Peptoclostridium difficile BI1 | methionyl-tRNA formyltransferase | *-1.568* | ***1.433*** |
| rpoB | CDBI1_00375 | Peptoclostridium difficile BI1 | DNA-directed RNA polymerase subunit beta | *-1.688* | ***1.428*** |
| CDBI1_04460 | CDBI1_04460 | Peptoclostridium difficile BI1 | hypothetical protein CDBI1_04460 | *-1.727* | ***1.315*** |
| CDBI1_00070 | CDBI1_00070 | Peptoclostridium difficile BI1 | seryl-tRNA synthetase | *-1.754* | ***1.787*** |

| | | | | | |
|---|---|---|---|---|---|
| CDBI1_13170 | CDBI1_13170 | Peptoclostridium difficile BI1 | amidohydrolase | *-1.781* | ***1.860*** |
| ileS | CDBI1_12715 | Peptoclostridium difficile BI1 | isoleucyl-tRNA synthetase | ***-1.847*** | ***1.563*** |
| CDBI1_01980 | CDBI1_01980 | Peptoclostridium difficile BI1 | putative fructose-bisphosphate aldolase | ***-1.953*** | ***3.021*** |
| tpiA | CDBI1_15495 | Peptoclostridium difficile BI1 | triosephosphate isomerase | ***-2.020*** | ***1.346*** |
| tsf | CDBI1_10370 | Peptoclostridium difficile BI1 | elongation factor Ts | ***-2.073*** | ***1.371*** |
| CDBI1_11280 | CDBI1_11280 | Peptoclostridium difficile BI1 | inosine 5-monophosphate dehydrogenase | ***-2.073*** | ***1.315*** |
| obgE | CDBI1_05225 | Peptoclostridium difficile BI1 | GTPase CgtA | ***-2.126*** | ***1.374*** |
| CDBI1_14055 | CDBI1_14055 | Peptoclostridium difficile BI1 | regulatory protease | ***-2.192*** | ***1.346*** |
| CDBI1_16190 | CDBI1_16190 | Peptoclostridium difficile BI1 | ATP-dependent protease La | ***-2.219*** | ***1.801*** |
| CDBI1_03635 | CDBI1_03635 | Peptoclostridium difficile BI1 | formate acetyltransferase | ***-2.272*** | ***1.401*** |
| CDBI1_08180 | CDBI1_08180 | Peptoclostridium difficile BI1 | guanine deaminase | ***-2.285*** | ***1.840*** |
| CDBI1_16485 | CDBI1_16485 | Peptoclostridium difficile BI1 | 6-phosphofructokinase | ***-2.299*** | ***1.518*** |
| CDBI1_11970 | CDBI1_11970 | Peptoclostridium difficile BI1 | heat shock protein | ***-2.312*** | ***1.600*** |
| QUI_3846 | QUI_3846 | Peptoclostridium difficile P59 | vitamin B12 dependent methionine synthase, activation domain protein | ***-2.352*** | ***1.338*** |
| CDBI1_13615 | CDBI1_13615 | Peptoclostridium difficile BI1 | S-layer precursor protein [Surface layer protein A] | ***-2.379*** | ***1.967*** |
| CDBI1_02215 | CDBI1_02215 | Peptoclostridium difficile BI1 | MarR family transcriptional regulator | ***-2.432*** | ***1.883*** |
| CDBI1_13610 | CDBI1_13610 | Peptoclostridium difficile BI1 | preprotein translocase subunit SecA | ***-2.432*** | ***1.754*** |
| CDBI1_12685 | CDBI1_12685 | Peptoclostridium difficile BI1 | peptidase | ***-2.525*** | ***2.259*** |
| CDBI1_09460 | CDBI1_09460 | Peptoclostridium difficile BI1 | hypothetical protein CDBI1_09460 [Coenzyme F420-0:L-glutamate ligase] | ***-2.631*** | ***2.625*** |
| CDBI1_01960 | CDBI1_01960 | Peptoclostridium difficile BI1 | acyl-CoA dehydrogenase family protein | ***-2.671*** | ***2.345*** |

| | | | | | |
|---|---|---|---|---|---|
| rplD | CDBI1_00415 | Peptoclostridium difficile BI1 | 50S ribosomal protein L4 | ***-2.697*** | ***1.373*** |
| cysS | CDBI1_00275 | Peptoclostridium difficile BI1 | cysteinyl-tRNA synthetase | ***-2.724*** | ***1.781*** |
| CDBI1_10585 | CDBI1_10585 | Peptoclostridium difficile BI1 | aromatic compounds hydrolase | ***-2.737*** | ***1.562*** |
| CDBI1_13425 | CDBI1_13425 | Peptoclostridium difficile BI1 | phosphoenolpyruvate-protein phosphotransferase | ***-2.777*** | ***1.987*** |
| CDBI1_01940 | CDBI1_01940 | Peptoclostridium difficile BI1 | isocaprenoyl-CoA:2-hydroxyisocaproate CoA-transferase | ***-2.804*** | ***1.342*** |
| aspS | CDBI1_13350 | Peptoclostridium difficile BI1 | aspartyl-tRNA synthetase | ***-2.870*** | ***1.349*** |
| CDBI1_01965 | CDBI1_01965 | Peptoclostridium difficile BI1 | electron transfer flavoprotein subunit beta | ***-2.883*** | ***2.352*** |
| CDBI1_01770 | CDBI1_01770 | Peptoclostridium difficile BI1 | putative aminopeptidase 2 | ***-2.883*** | ***1.643*** |
| CDBI1_13210 | CDBI1_13210 | Peptoclostridium difficile BI1 | cell surface protein [Cell wall binding protein Cwp22] | ***-2.897*** | ***1.349*** |
| CDBI1_13280 | CDBI1_13280 | Peptoclostridium difficile BI1 | putative serine hydroxymethyltransferase | ***-2.937*** | ***2.908*** |
| CDBI1_04210 | CDBI1_04210 | Peptoclostridium difficile BI1 | ABC transporter substrate-binding protein | ***-2.937*** | ***1.846*** |
| CDBI1_13585 | CDBI1_13585 | Peptoclostridium difficile BI1 | cell surface-associated cysteine protease [Cell wall binding protein Cwp84] | ***-2.937*** | ***1.846*** |
| CDBI1_16105 | CDBI1_16105 | Peptoclostridium difficile BI1 | glucose-6-phosphate isomerase | ***-3.030*** | ***2.258*** |
| CDBI1_11210 | CDBI1_11210 | Peptoclostridium difficile BI1 | transketolase | *-3.056* | ***3.527*** |
| CDBI1_13050 | CDBI1_13050 | Peptoclostridium difficile BI1 | pyruvate-flavodoxin oxidoreductase | ***-3.069*** | ***2.510*** |
| CDBI1_16960 | CDBI1_16960 | Peptoclostridium difficile BI1 | F0F1 ATP synthase subunit beta | ***-3.176*** | ***2.746*** |
| CDBI1_15505 | CDBI1_15505 | Peptoclostridium difficile BI1 | glyceraldehyde-3-phosphate dehydrogenase 2 | ***-3.176*** | ***2.528*** |
| tig | CDBI1_16215 | Peptoclostridium difficile BI1 | trigger factor | ***-3.176*** | ***1.358*** |
| CdifQCD-_020200013241 | CdifQCD-_020200013241 | Clostridium difficile QCD-97b34 | pyruvate-flavodoxin oxidoreductase | ***-3.242*** | ***2.018*** |

| | | | | | |
|---|---|---|---|---|---|
| rplK | CDBI1_00350 | Peptoclostridium difficile BI1 | 50S ribosomal protein L11 | ***-3.269*** | ***1.470*** |
| CDBI1_00930 | CDBI1_00930 | Peptoclostridium difficile BI1 | redox-sensing transcriptional repressor Rex | ***-3.295*** | ***2.421*** |
| CDBI1_03375 | CDBI1_03375 | Peptoclostridium difficile BI1 | aminoacyl-histidine dipeptidase | ***-3.415*** | ***1.998*** |
| CDBI1_00265 | CDBI1_00265 | Peptoclostridium difficile BI1 | prolyl-tRNA synthetase | ***-3.442*** | ***2.961*** |
| thiH | CDBI1_10450 | Peptoclostridium difficile BI1 | thiamine biosynthesis protein ThiH | ***-3.468*** | ***1.650*** |
| greA | CDBI1_17400 | Peptoclostridium difficile BI1 | transcription elongation factor GreA | ***-3.614*** | ***2.091*** |
| CDBI1_03405 | CDBI1_03405 | Peptoclostridium difficile BI1 | adenosylcobamide-dependent radical SAM protein | ***-3.628*** | ***1.344*** |
| CDBI1_01505 | CDBI1_01505 | Peptoclostridium difficile BI1 | hypothetical protein CDBI1_01505 | ***-3.760*** | ***1.590*** |
| CDBI1_16090 | CDBI1_16090 | Peptoclostridium difficile BI1 | formate acetyltransferase | ***-3.960*** | ***1.373*** |
| CDBI1_13380 | CDBI1_13380 | Peptoclostridium difficile BI1 | adenine phosphoribosyltransferase | ***-4.053*** | ***1.476*** |
| rplF | CDBI1_00485 | Peptoclostridium difficile BI1 | 50S ribosomal protein L6 | ***-4.265*** | ***2.321*** |
| CDBI1_17105 | CDBI1_17105 | Peptoclostridium difficile BI1 | DNA-binding protein HU | ***-4.823*** | ***2.403*** |
| CDBI1_19448 | CDBI1_19448 | Peptoclostridium difficile BI1 | hypothetical protein CDBI1_19448 | ***-6.298*** | ***1.328*** |
| CDBI1_05205 | CDBI1_05205 | Peptoclostridium difficile BI1 | ribonuclease g | ***-6.405*** | ***1.400*** |
| CDBI1_00450 | CDBI1_00450 | Peptoclostridium difficile BI1 | 50S ribosomal protein L29 | ***-6.431*** | ***1.509*** |
| CDBI1_01190 | CDBI1_01190 | Peptoclostridium difficile BI1 | dtdp-4-dehydrorhamnose reductase | ***-6.498*** | ***1.484*** |

Red indicates high abundance; green indicates less abundance. Increasing fold changes are reported as brighter color.
Fold changes reported in Log2 values.
P-values reported as -log10 values.

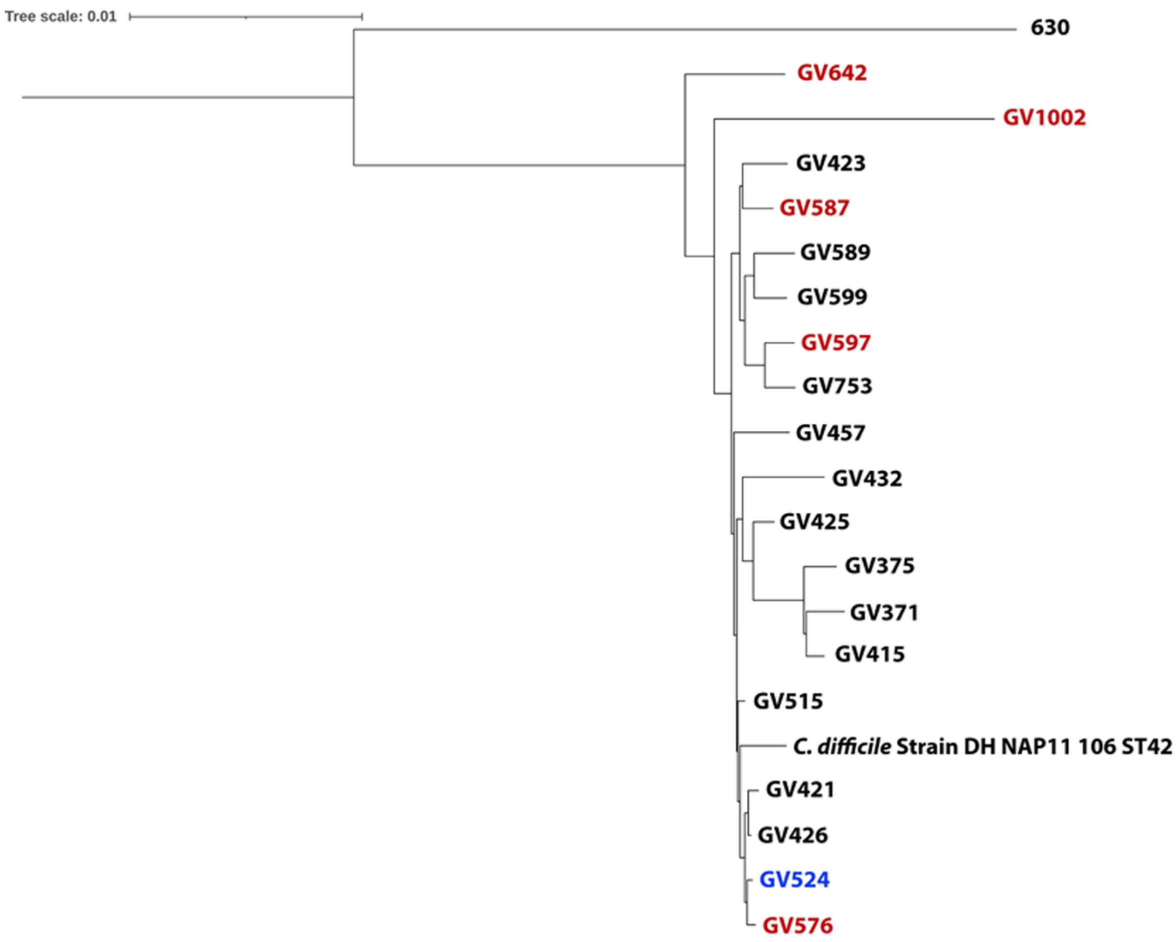


**Supplemental Figure S3: Composite Vector Tree of High- and Low-Toxin RT106 Strains**

Whole genome comparison was done between clinically derived, low toxin producing (n = 5; red) and high toxin producing strains (n = 14; black) using CVTree3 (http://tlife.fudan.edu.cn/cvtree3/). Strain 630 is a low toxin comparator and GV524 is a Tox$^-$/NAAT$^-$ strain. Whole genome comparison of these strains show that low-toxin strains do not clade separately and is interspersed between high-toxin strains.

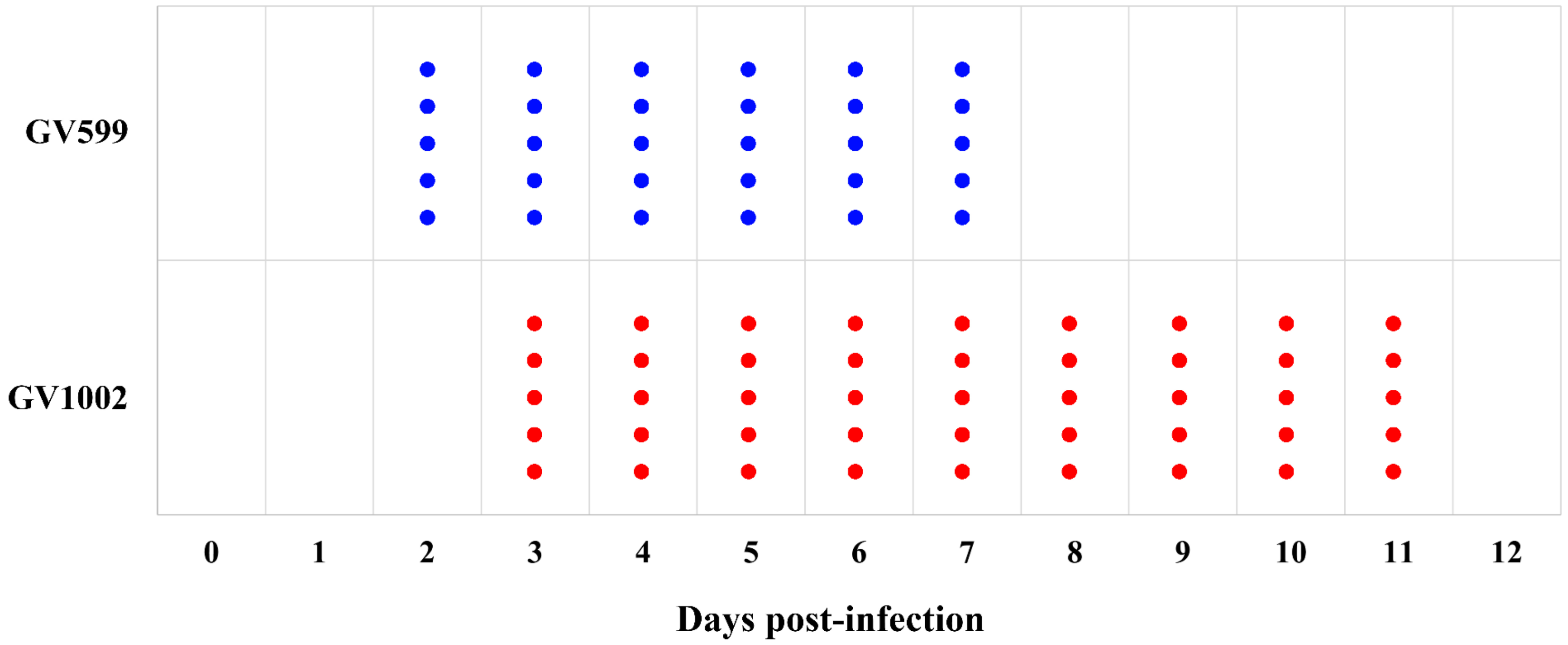


**Supplemental Figure S4: Mouse Infection with High- and Low-toxin RT106 Strains**

C57L/6 mice (n = 5 / group) were infected with a high-toxin RT106 strain, GV599 (blue), or a low-toxin RT106 strain, GV1002 (red). Each dot represents an animal that we have detected *C. difficile* in the stool. The high-toxin strain GV599 was detected on Day 2 post-infection and subsequently cleared in all mice by Day 8 post-infection resulting in a 6-day infection. The low-toxin strain was detected on Day 3 post-infection and was subsequently cleared by all mice by Day 12 post-infection resulting in a 9-day infection.

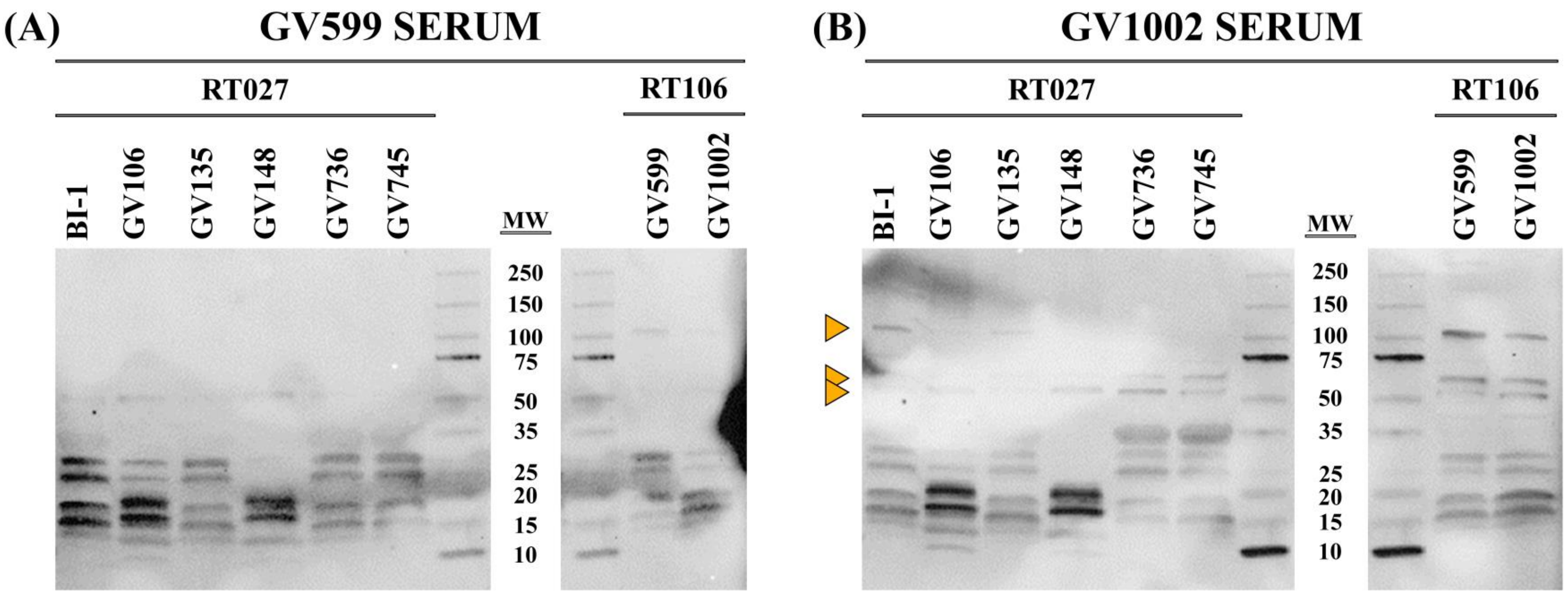


**<u>Supplemental Figure S5: Membrane Fractions Probed with High- and Low-toxin RT106 Infected Mouse Serum</u>**

Membrane extracts from high-toxin strains BI-1 (RT027) and GV599 (RT106), as well as the low-toxin strains GV106, GV135, GV148, GV736, GV745 (RT027s) and GV1002 (RT106) were probed with serum from mice infected with GV599 and GV1002. As with probing with RT027-infected mouse serum, the low-toxin strain serum (GV1002) bound to distinct bands that were not found when probing with high-toxin strain serum (GV599). These bands are indicated with orange triangles and were found in all membrane fractions suggesting an altered cell surface of the low-toxin strains irrespective of ribotypes.

### Supplemental Table S8: Discrepant RT106-specific Genes Compared to Discrepant RT027 Genes.

| Gene ID | AA Length | Function | Found In LT027 Pangenome? | Found in GV148? | Found in BI-1? |
|---|---|---|---|---|---|
| fig\|1496.6344.peg.314 | 913 | PAS domain histidine kinase | N | N | N |
| fig\|1496.6344.peg.709 | 187 | Mobile element protein | N | N | N |
| fig\|1496.6344.peg.831 | 98 | Mobile element protein | N | N | N |
| fig\|1496.6344.peg.1058 | 308 | hypothetical protein | N | Y | Y |
| fig\|1496.6344.peg.1149 | 1129 | Transcription-repair coupling factor | N | Y | Y |
| fig\|1496.6344.peg.1255 | 123 | Functional domain: RNA-binding protein YhbY | Y | Y | N |
| fig\|1496.6344.peg.1360 | 39 | hypothetical protein | N | N | N |
| fig\|1496.6344.peg.1711 | 458 | Arginine utilization regulatory protein RocR | N | Y | Y |
| fig\|1496.6344.peg.1854 | 44 | LytR family transcriptional regulator domain | N | N | N |
| fig\|1496.6344.peg.1937 | 200 | hypothetical protein | N | Y | Y |
| fig\|1496.6344.peg.2047 | 898 | Multimodular transpeptidase-transglycosylase (EC 2.4.1.129) (EC 3.4.-.-) | N | Y | N |
| fig\|1496.6344.peg.2128 | 637 | Phosphonoacetaldehyde hydrolase (EC 3.11.1.1) / 2-aminoethylphosphonate:pyruvate aminotransferase (EC 2.6.1.37) | N | Y | Y |
| fig\|1496.6344.peg.2343 | 50 | hypothetical protein | N | N | N |
| fig\|1496.6344.peg.2544 | 289 | FIG00512787: hypothetical protein | N | Y | Y |
| fig\|1496.6344.peg.2729 | 445 | Pyridine nucleotide-disulfide oxidoreductase; NADH dehydrogenase (EC 1.6.99.3) | N | Y | Y |
| fig\|1496.6344.peg.2831 | 44 | hypothetical protein | N | N | N |
| fig\|1496.6344.peg.3114 | 699 | CocE/NonD family hydrolase | N | Y | Y |
| fig\|1496.6344.peg.3564 | 425 | Two-component sensor histidine kinase | N | Y | Y |

Discrepant RT106 strains (N=5); non-discrepant RT106 strains (N=4)
